# Strain-level differences in gut microbiome composition determine fecal IgA levels and are modifiable by gut microbiota manipulation

**DOI:** 10.1101/544015

**Authors:** Chao Yang, Ilaria Mogno, Eduardo J. Contijoch, Joshua N. Borgerding, Varun Aggarwala, Sophia Y. Siu, Zhihua Li, Emilie K. Grasset, Drew S. Helmus, Marla C. Dubinsky, Saurabh Mehandru, Andrea Cerutti, Jeremiah J. Faith

**Author notes:** Correspondence (J.J.F.).

## Abstract

Fecal IgA production depends on colonization by a gut microbiota. However, the bacterial strains that drive gut IgA production remain largely unknown. By accessing the IgA-inducing capacity of a diverse set of human gut microbial strains, we identified *Bacteroides ovatus* as the species that best induced gut IgA production. However, this induction varied bimodally across different *B. ovatus* strains. The high IgA-inducing *B. ovatus* strains preferentially elicited more IgA production in the large intestine largely through the T-cell-dependent B cell-activation pathway. Remarkably, a low-IgA phenotype in mice could be robustly and consistently converted into a high-IgA phenotype by transplanting a multiplex cocktail of high IgA-inducing *B. ovatus* strains but not individual ones. Our results highlight the critical importance of microbial strains in driving phenotype variation in the mucosal immune system and provide a strategy to robustly modify a gut immune phenotype, including IgA production.

## Introduction

Immunoglobulin A (IgA) is the most abundant mucosal antibody and plays an essential role in maintaining gut homeostasis as well as other physiological processes (Cerutti and Rescigno, 2008; Macpherson et al., 2012; Sutherland et al., 2016). Secretory IgA, for example, can limit the access of bacteria and bacteria-derived toxins to intestinal epithelial cells (Okai et al., 2016; Tokuhara et al., 2010), facilitate the clearance of bacteria that have breached the mucosal barrier (Fagarasan, 2008; Pabst, 2012; Strugnell and Wijburg, 2010) and regulate the colonization of bacteria in the mucosal lining (Donaldson et al., 2018; McLoughlin et al., 2016). In addition, IgA can also bind disease-associated gut microbiota (Kau et al., 2015; Palm et al., 2014; Viladomiu et al., 2017). Conversely, the gut microbiota and its metabolites drive the production of IgA as germ-free (GF) mice have an almost undetectable level of fecal IgA (Kim et al., 2016; Macpherson et al., 2000). Upon bacteria colonization, even with a single bacterial strain (Fritz et al., 2011; Hapfelmeier et al., 2010; Peterson et al., 2007), B cells undergo class-switch to IgA^+^ cells in gut-associated lymphoid tissues (GALT), which include Peyer’s patches (PP), isolated lymphoid follicles (ILF) and mesenteric lymph nodes (MLN), and in the gut lamina propria (LP) (Macpherson et al., 2008; Pabst, 2012). Much of the intestinal IgA is bacteria-specific (Bunker et al., 2015; Hapfelmeier et al., 2010; Peterson et al., 2007), and the B-cell repertoire is highly influenced by the microbiota composition (Lindner et al., 2015). To date, a few murine derived bacterial species have been identified as being able to enhance or reduce intestinal IgA level (Chudnovskiy et al., 2016; Lecuyer et al., 2014; Moon et al., 2015; Obata et al., 2010). However, key questions regarding the impact of microbiota in this process remain largely unanswered including the importance of colonization order, the contribution of individual bacterial species versus that of microbial communities, the potential to modulate IgA production by altering gut microbiota composition with commensal organisms, and the role of each microbial species in the development of IgA^+^ B cells in specific tissues (Macpherson et al., 2015; Pabst, 2012).

Apart from IgA-secreting cells, the gut microbiota has the capacity to influence numerous other immune cell populations including colonic regulatory T cells (Treg) (Atarashi et al., 2011; Faith et al., 2014; Round and Mazmanian, 2010), IL-17 producing T helper cells (Ivanov et al., 2009), and macrophages (Mortha et al., 2014). Importantly, many of these responses seem to be bacterial strain-specific as communities with comparable species composition can drive gut immune responses characterized by largely different cell compositions (Britton et al., 2019). These discoveries indicate that manipulation of the gut microbiota, with appropriate bacterial strains, represents a potential therapeutic pathway for the treatment of diseases including inflammatory bowel disease (IBD), rheumatoid arthritis (RA) and multiple sclerosis through shaping the host immune system (Skelly et al., 2019). Although the studies of microbiota-based therapeutics and fecal microbiota transplantation (FMT) have heavily focused on the engraftment of the transmitted microbiota and its influence on the composition of the recipient microbiota (Seekatz et al., 2014; Shankar et al., 2014; Smillie et al., 2018), the clinical application of microbiota manipulation as an immunomodulatory strategy will require combinations of bacterial strains optimized for the induction of specific immune phenotypes that are robust to the interpersonal variation in the pre-existing microbiota of each recipient.

Here we demonstrate that, upon transfer into GF mice, human isolates of the *Bacteroides ovatus* species, one of the most common human gut commensals, are uniquely capable of inducing high mucosal IgA production compared with other common gut commensal species. This IgA-inducing capacity, however, was restricted to specific strains of *B. ovatus* that preferentially led to IgA production in the large intestine through both T-cell-dependent (TD) and T-cell-independent (TI) B cell-activation pathways. While no individual bacterial strain functioned as an effective enhancer of gut IgA production, we found that cocktails of these high IgA-inducing (IgA^high^) strains could serve as effective immunomodulators, that elicited higher fecal IgA levels upon administration to animals harboring a pre-existing microbiota with low IgA-inducing potential (IgA^low^). Our work demonstrates the importance of strain-level variation in gut microbiota composition on mucosal immune responses. It also supports the potential utility of cultured multi-bacterial effector strain cocktails as a strategy to overcome phenotype transfer resistance in microbiota-based immunomodulation (Petrof and Khoruts, 2014).

## Results

### *B. ovatus* elicits robust gut IgA production

To determine if individual gut bacterial species have a distinct IgA-inducing potential, we monocolonized GF C57BL/6 mice with one of eight different human gut commensal bacteria (Table S1) with representatives from the most prominent phyla of the human gut including Firmicutes, Bacteroidetes, Actinobacteria and Proteobacteria (Human Microbiome Project, 2012; Turnbaugh et al., 2009). After three weeks of colonization to allow optimal steady-state gut IgA secretion (Figure S1A), we measured serum and fecal IgA levels in each group of gnotobiotic mice (Peterson et al., 2007). Although all tested species significantly increased IgA level relative to control GF mice, *B. ovatus* monocolonized mice secreted significantly more IgA in their feces compared with mice colonized with any of the other seven human gut bacteria (Figure 1A; *p* < 0.001). Most species also increased serum IgA (Figure S1B). However, consistent with previous reports (Macpherson et al., 2008), fecal IgA and serum IgA levels in these mice did not correlate significantly (Figure S1C; R^2^ = 0.226; *p* = 0.196). GF mice colonized with the cocktail of all eight bacterial species yielded as much fecal and serum IgA as mice monocolonized with *B. ovatus*.

**Figure 1.**
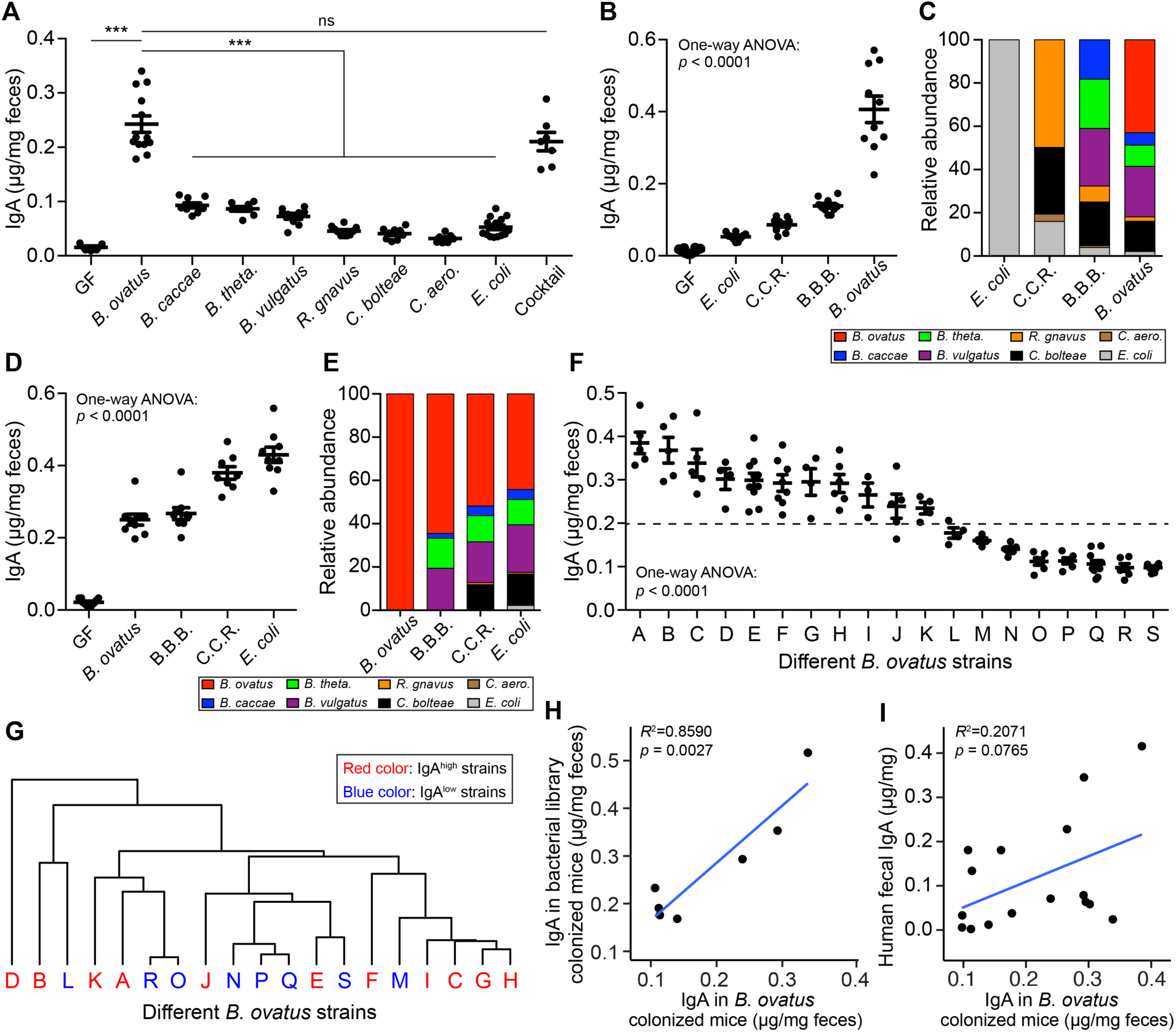
*B. ovatus* species, with strain-level differences, predominantly induces fecal IgA production in gnotobiotic mice. (**A**) Fecal IgA level in C57BL/6 gnotobiotic mice colonized with individual or a cocktail of human gut commensal bacteria for three weeks. (**B-E**) The concentration of fecal IgA (**B** and **D**) and proportion of each bacterial strain (**C** and **E**) in stool of gnotobiotic mice that were colonized sequentially with individual or combined bacterial communities starting from *E. coli* (**B** and **C**) or *B. ovatus* (**D** and **E**). Feces were harvested before addition of new bacteria to the same mice. C.C.R.: cocktail of *C. bolteae, C. aerofaciens* and *R. gnavus*; B.B.B.: cocktail of *B. caccae, B. theta*. and *B. vulgatus* (**F**) Quantification of fecal IgA in gnotobiotic mice upon colonization with an individual strain of *B. ovatus* for three weeks. Unique strains of *B. ovatus* were isolated from the stools of different human donors. Dotted line separates high- and low-IgA phenotypes. (**G**) Dendrogram clustering of different *B. ovatus* strains basing on the dissimilarity of bacterial genomic DNA sequences. (**H**) Correlation of stool IgA levels between single *B. ovatus* strain monocolonized mice versus mice colonized with a microbiota arrayed culture collection that included that single *B. ovatus* strain. Both single *B. ovatus* strains and arrayed culture collections were isolated from the same donor. (**i**) Correlation of fecal IgA concentration between single *B. ovatus* strain colonized mice versus human donor. Data shown are mean ± standard error of the mean. Each dot in **A**, **B**, **D** and **F** represents a biological replicate. The average fecal IgA concentration from 4-10 mice was used for correlation in **H** and **I**. Detailed strain information is listed in Tables S1 and S2. *p*-values with statistical significance (assessed by two-tailed Student’s *t* test or one-way ANOVA) are indicated: \*\*\**p* < 0.001; ns, not significant.

To address if the order of bacterial colonization could influence fecal IgA secretion, GF mice were sequentially colonized every three weeks with individual species or small cocktails of the same eight bacterial species. We first assayed fecal IgA level in mice sequentially colonized with low IgA inducers (i.e. *E. coli*) to high IgA inducers (i.e. *B. ovatus*) (Figure S1D). Fecal IgA increased gradually with the colonization of additional bacterial species. However, the more striking (>2-fold) increase in IgA occurred after colonization with *B. ovatus* (Figure 1B). Metagenomic sequencing of fecal microbiota in these mice revealed gut colonization by each bacterial species, albeit with different proportions (Figure 1C). We then reversed the order of colonization from high IgA inducers (i.e. *B. ovatus*) to low IgA inducers (i.e. *E. coli*) (Figure S1D). Once again, *B. ovatus* elicited the largest increase of fecal IgA production, while the other species led to smaller increases (Figure 1D). Remarkably, the relative abundance of each organism at the end of the colonization was very similar, regardless of the order of colonization (Figures 1C and 1E). These results demonstrate that *B. ovatus* is a uniquely potent gut IgA inducer and that the species composition of the gut microbiota impacts IgA production more than the order of bacterial colonization.

To test the role of bacterial viability in the induction of gut IgA by *B. ovatus* (Hapfelmeier et al., 2010), GF mice were administered heat-killed *B. ovatus* or *B. ovatus* metabolites (i.e. filtered, conditioned growth medium from stationary phase of *B. ovatus* cultures) for three weeks. Neither approach was capable of enhancing fecal IgA above the level detected in GF mice (Figure S1E). To ensure the above result was not due to the underdeveloped mucosal immune system of GF mice, we performed similar experiments by first colonizing GF mice with *E. coli* for three weeks and subsequently treated these mice with heat-killed *B. ovatus* for an additional three weeks. Again, we found no significant fecal IgA increase (Figure S1E). Thus, neither dead *B. ovatus* nor its metabolites triggered efficient gut IgA responses in the murine intestine. All together, live *B. ovatus* species elicited more gut IgA production than other tested gut commensal bacterial species in GF mice.

### *B. ovatus*-driven gut IgA production is strain-specific

Given the remarkable microbial strain variation across individuals (Faith et al., 2015; Faith et al., 2013; Vatanen et al., 2018; Zhao et al., 2019), we wondered whether all *B. ovatus* strains within this common bacterial species induced comparably high fecal IgA. GF mice monocolonized for three weeks with one of 19 *B. ovatus* strains isolated from 19 different individuals (Table S2) showed a strain-specific gut IgA response (Figure 1F; *p* < 0.0001). In contrast to the large variability of fecal IgA levels, serum IgA levels were comparable across mice monocolonized with different *B. ovatus* strains (Figure S2A). Similarly, the colonization density was also comparable across mice harboring different *B. ovatus* strains (Figure S2B). This observation suggests that the global density of each individual strain was not implicated in the genesis of strain-specific differences of gut IgA responses. Of note, the distribution pattern of IgA induction across multiple *B. ovatus* strains was bimodal (Figure S2C; *p* = 0.0481 Hartigans’ Dip Test), allowing these strains to be categorized as IgA^high^ or IgA^low^. The genomic similarity of *B. ovatus* strains was not a significant predictor of their IgA^high^ and IgA^low^ properties (Figure 1G and Table S3), which suggests that their distinct IgA-inducing function is shared amongst the species rather than representing an evolutionarily distinct group within the species.

To rule out a bias in our preliminary screen for IgA^low^ strains within the *Bacteroides* genus, we assayed whether additional strains could induce high fecal IgA (Figure 1A). We found no strain-specific differences in fecal IgA induction when GF mice were monocolonized with three distinct strains of *B. caccae*, *B. thetaiotaomicron* and *B. vulgatus* (Figure S2D). The IgA-inducing function of additional common species from the order Bacteroidales, including *Parabacteroides johnsonii*, *Bacteroides intestinalis* and *Bacteroides fragilis*, were tested but also induced much less gut IgA than *B. ovatus* (Figure S2D). These results indicate that the high IgA-inducing ability of *B. ovatus* is unique to this gut bacterial species and only to a subset of strains.

To examine the influence of *B. ovatus* strain variation on host fecal IgA production in the context of more complex gut microbiotas, we colonized GF mice with one of the seven microbiota arrayed culture collections originally isolated from different human donors with each collection consisting of 15-20 unique species (Britton et al., 2019). The arrayed culture collections were assembled to reconstitute a donor microbiota each containing a unique *B. ovatus* strain, which was already functionally tested by earlier monocolonization (Figure 1F). We observed a significant positive correlation between the fecal IgA concentrations induced by an individual *B. ovatus* strain and the fecal IgA concentrations elicited by a culture collection representing the entire *B. ovatus*-containing microbiota from the same donor (Figures 1H and S2E; R^2^ = 0.859, *p* = 0.0027). Again, these results suggest that the *B. ovatus* strain composition is a major contributor of gut IgA responses even when considered in the context of complex microbial communities.

Unlike inbred laboratory mice housed in a highly controlled environment, human beings, with different genetic background, are exposed to more complex continuum of factors including some that were demonstrated to affect fecal IgA production such as genetics and diet (Fransen et al., 2015; Kim et al., 2016). To determine whether *B. ovatus* could drive robust gut IgA responses also in humans, we measured the fecal concentration of IgA in multiple human donors and correlated this concentration with that of fecal IgA generated by GF mice monocolonized with a *B. ovatus* strain isolated from identical donors. Though no significant correlation was observed, there was a clear trend towards a positive correlation even in an uncontrolled condition (Figures 1I and S2E; R^2^ = 0.2071, *p* = 0.0765).

In total these results demonstrate that a subset of *B. ovatus* strains induce high fecal IgA levels, which broadly influence the total fecal IgA output of the host even in the context of a diverse gut microbiota.

### IgA^high^ *B. ovatus* strains induce more IgA production in the large intestine

To interrogate the mechanisms underpinning gut IgA induction by different *B. ovatus* strains, GF mice were colonized with a representative IgA^high^ or IgA^low^ strain (*B. ovatus* strain E and Q, respectively). We quantified bacteria-bound IgA in the stool of mice. Monocolonization with the IgA^high^ strain E not only induced more free fecal IgA but also more fecal bacteria-bound IgA than the IgA^low^ strain Q (52.9% vs. 21.0% IgA-coated *B. ovatus*) (Figure 2A). In contrast, no significant difference was observed in serum immunoglobulin isotypes (i.e. IgA, IgG1, IgG2a, IgG2b, IgG3, IgM and IgE) in monocolonized mice harboring either *B. ovatus* strain E or Q (Figures S2A and S3A).

**Figure 2.**
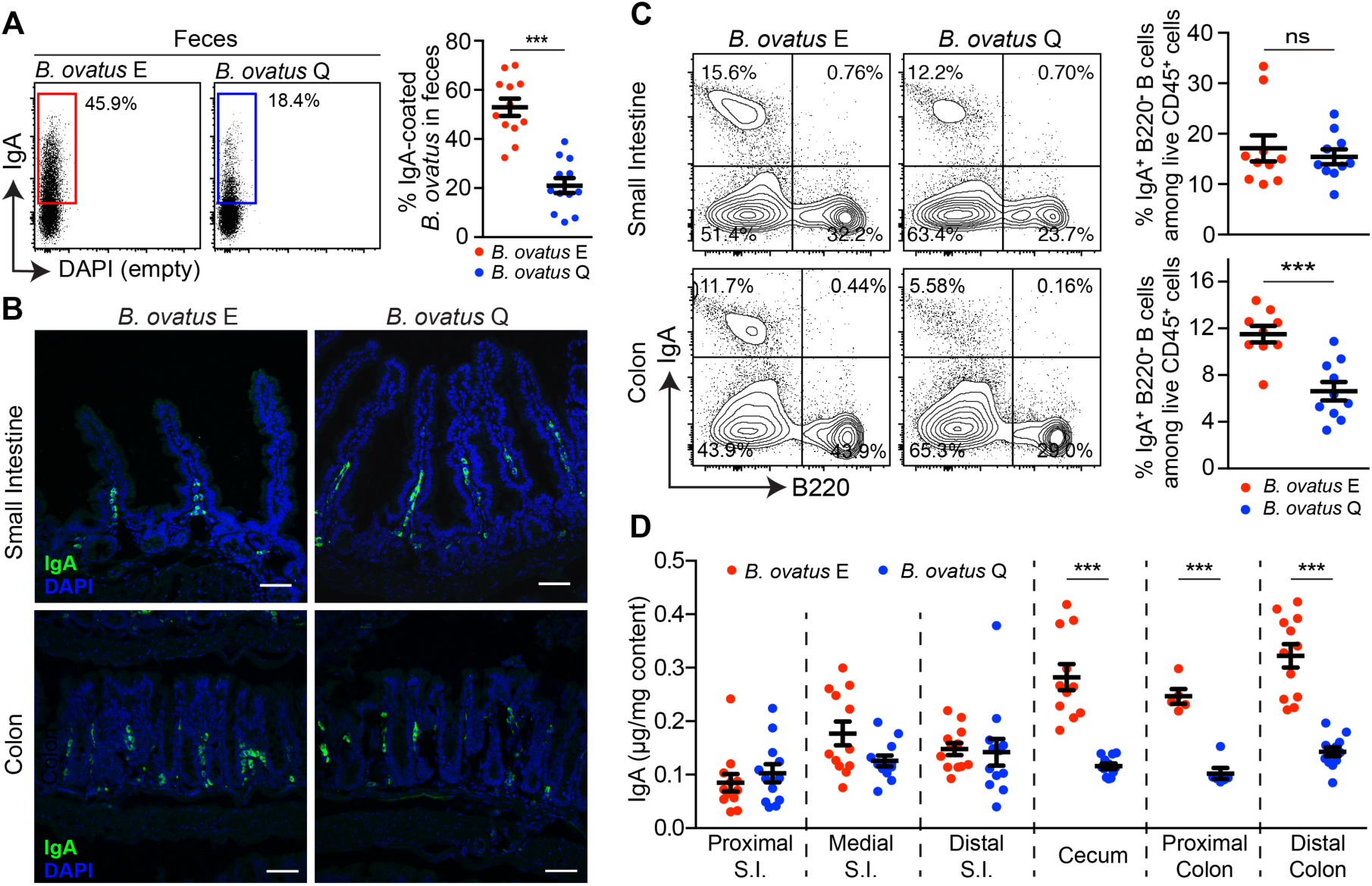
IgA^high^ *B. ovatus* strain Elicits stronger IgA responses in the large intestine. (**A**) Representative flow cytometry plot and quantification of IgA-coated *B. ovatus* in feces of gnotobiotic mice harboring either *B. ovatus* strain E or Q. (**B**) Representative images of IgA^+^ cells in small intestine and the colon are shown. IgA^+^ cells were stained with anti-IgA (green); Nuclei were counter-stained with DAPI (4’,6-diamidino-2-phenylindole) (blue). n = 5~6. Scale bar = 50 μm. (**C**) Percentage of IgA^+^ B cells, analyzed by flow cytometry, in small intestine and colon is shown. Number adjacent to gate represents percentage. (**D**) Free IgA concentration in luminal contents along the length of the intestine. S.I.: small intestine. Data shown are mean ± standard error of the mean. Each dot represents an individual mouse. *p*-values with statistical significance (assessed by two-tailed Student’s *t* test) are indicated: \*\*\**p* < 0.001; ns, not significant.

Fecal IgA mostly derives from polymeric IgA released by IgA^+^ plasma cells residing in the intestinal LP and translocated to the gut lumen across epithelial cells via transcytosis (Johansen and Kaetzel, 2011). This process is mediated by a basolateral IgA (and IgM) transporter termed polymeric immunoglobulin receptor (pIgR) (Johansen and Kaetzel, 2011). Independent groups have reported that the expression of pIgR by gut epithelial cells is influenced by bacteria stimulation both *in vivo* and *in vitro* (Hooper et al., 2001; Schneeman et al., 2005). To determine if *B. ovatus* strain variation impacts fecal IgA level by modulating pIgR-mediated transcytosis, we imaged the expression of pIgR by immunofluorescence staining in the small intestine and the colon of mice colonized with either *B. ovatus* strain E (IgA^high^) or Q (IgA^low^). However, no noticeable difference in protein expression or mRNA transcription of pIgR was observed (Figures S3B and S3C). We also found that the two strains of *B. ovatus* colonized the colon similarly without penetrating into epithelial cells and induced similar level of mRNA transcription of Muc2 (Figures S3C and S3D). To further interrogate the mechanism underpinning the increased fecal IgA in *B. ovatus* strain E colonized mice, we then quantified, by histology and flow cytometry, IgA-secreting B cells in both small intestine and the colon. We found more IgA-secreting B cells in the colonic LP of mice harboring *B. ovatus* strain E compared to mice harboring strain Q, while no significant strain-specific difference was observed in the small intestine (Figures 2B and 2C). Although PPs and MLNs usually serve as dominant IgA inductive sites (Chorny et al., 2010; Fagarasan et al., 2010), we did not find a significant difference in IgA^+^ B cells induction at these sites by strain E or Q (Figures S4A and S4B).

Given the preferential expansion of IgA-secreting B cells in the colon of monocolonized mice harboring *B. ovatus* strain E, we then explored whether luminal IgA levels would vary between small and large intestinal regions. In the small intestine, we found that mice monocolonized with *B. ovatus* strain E or Q had comparable luminal IgA levels (Figure 2D). In contrast, mice monocolonized with strain E had significantly more luminal IgA from cecum to distal colon than those colonized with strain Q (Figure 2D). Similar results were also observed across all tested IgA^high^ and IgA^low^ *B. ovatus* strains (Figure S5). Thus, the IgA^high^ *B. ovatus* strains induce more colonic IgA-secreting B cells compared to IgA^low^ *B. ovatus* strains, which results in the secretion of more IgA to the large intestinal lumen.

To determine if these observations were unique to GF C57BL/6 mice, we recapitulated our monocolonization strategy in GF Swiss Webster mice and found that fecal IgA was largely comparable in gnotobiotic C57BL/6 and Swiss Webster mice colonized with identical bacterial strains (Figures S6A-S6D; R^2^ = 0.601, *p* = 0.0011). Moreover, IgA^high^ strain colonized gnotobiotic Swiss Webster mice also secreted more intraluminal IgA in the large intestine compared with IgA^low^ strain colonized mice (Figure S6E). Thus, bacteria-induced gut IgA production is similar across different host genetic backgrounds.

### *B. ovatus* elicits gut IgA production primarily via TD B cell-activation pathway

Gut IgA responses occur through TD and/or TI B cell-activation pathways (Fagarasan et al., 2010). To determine the influence of CD4^+^ T cells on the gut IgA production induced by *B. ovatus*, we depleted CD4^+^ T cells in mice by injecting with an anti-CD4 antibody five days prior to and for three weeks after monocolonization with *B. ovatus* strain E or Q (Figures 3A and S7A-S7C). On day seven post-colonization, fecal IgA increased in both T cell-depleted and T cell-sufficient gnotobiotic mice, which suggests that CD4^+^ T cells are not a dominant factor in early stage IgA induction. By day 14 post-colonization, control mice receiving an isotype-matched irrelevant antibody generated significantly more fecal IgA than mice receiving anti-CD4 antibody suggesting the majority of the steady state *B. ovatus* induced IgA is T cell dependent (Figure 3B). In addition to reduced free IgA, *B. ovatus*-bound IgA also decreased in the stool of CD4^+^ T cell-depleted mice (Figure 3C) in both *B. ovatus* strain E or Q colonized mice. Moreover, in the small intestine and the colon, the frequency of IgA-secreting B cells was reduced significantly compared to that of IgA-secreting B cells being detected in the control CD4^+^ T cell-sufficient mice (Figures 3D and S7D-S7G). In addition, these control mice showed more intraluminal IgA than CD4^+^ T cell-depleted mice across the whole intestinal tract (Figure 3E).

**Figure 3.**
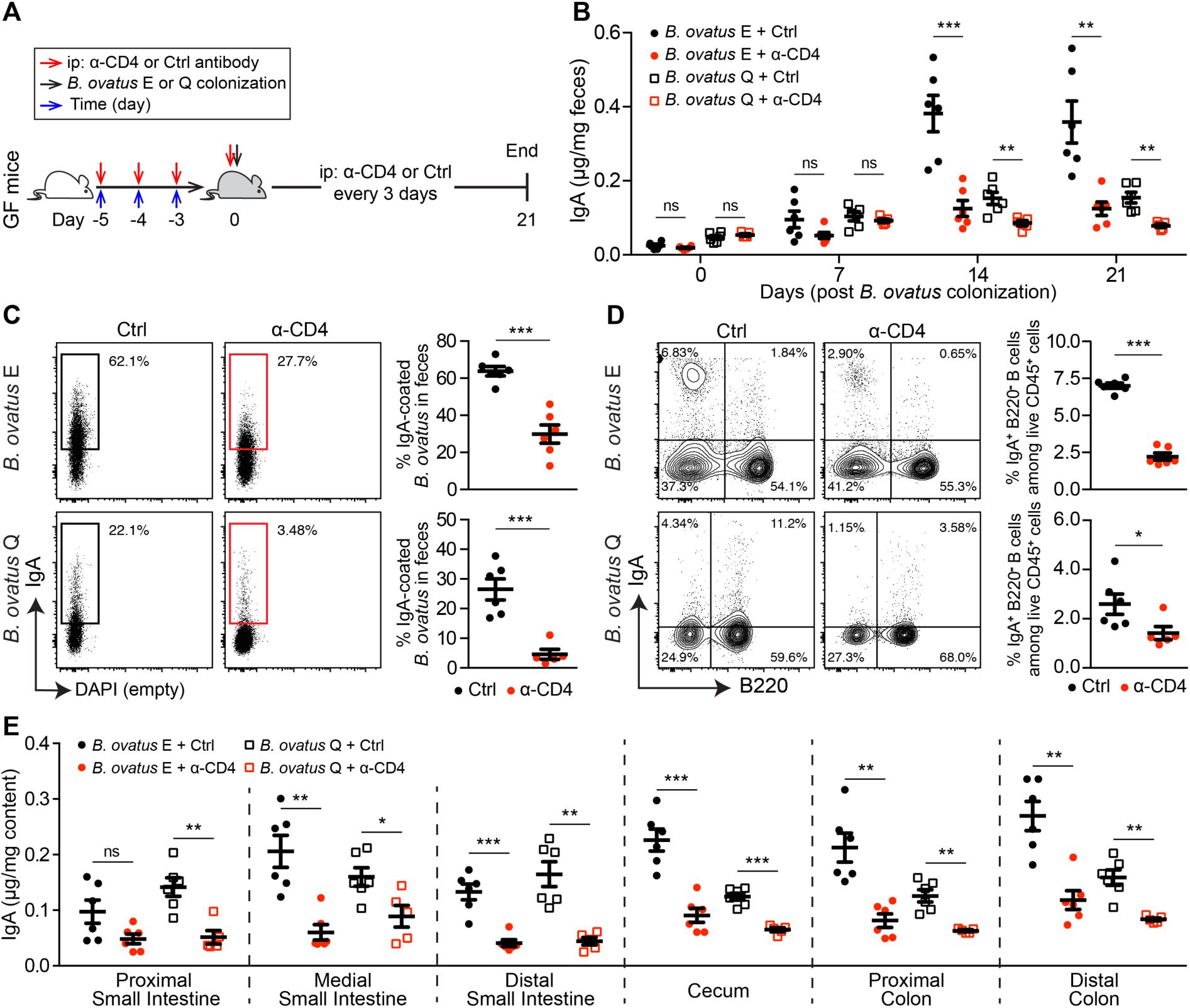
T-cell-dependent B cell activation pathway plays an essential role in *B. ovatus* induced fecal IgA production. (**A**) Schematic representation of CD4^+^ T cells depletion in germ-free B6 mice is illustrated. Red arrows represent i.p. injection of anti-CD4 antibody or isotype control. Black arrow indicates *B. ovatus* strain E or Q colonization and blue arrows represent time. (**B**) Dynamics of fecal IgA concentration in *B. ovatus* strain E or Q colonized gnotobiotic B6 mice treated with either anti-CD4 antibody or isotype control. (**C**) Representative flow cytometry plot and quantification of IgA-coated bacteria in feces of *B. ovatus* strain E or Q colonized gnotobiotic mice treated with either anti-CD4 antibody or isotype control. (**D**) Representative flow cytometry plot and percentage of IgA-secreting B cells in the colon of mice colonized with *B. ovatus* strain E or Q w/o anti-CD4 antibody treatment are shown. Numbers adjacent to gates represent percentage. (**E**) Concentration of free IgA in the intestinal content collected from different regions of the whole intestinal tract is shown. Data shown are mean ± standard error of the mean. Each dot represents an individual mouse. *p*-values with statistical significance (assessed by two-tailed Student’s *t* test) are indicated: \**p* < 0.05, \*\**p* < 0.01, \*\*\**p* < 0.001; ns, not significant.

### Multiplex cocktail of *B. ovatus* strains robustly modify gut IgA production

Given the potential of gut microbiota manipulation as a therapeutic, we next determined whether the high-IgA phenotype could be transferred to mice harboring microbiotas that induce a low level of fecal IgA. For this purpose, we recolonized GF C57BL/6 mice with either *B. ovatus* strain E (IgA^high^) or Q (IgA^low^) for three weeks, followed by cohousing these mice for an additional three weeks (Figure 4A). After cohousing, mice monocolonized with *B. ovatus* strain Q showed no significant change in fecal IgA. In contrast, mice colonized initially with *B. ovatus* strain E had reduced fecal IgA, which raised the possibility that the low-IgA phenotype behaves as a dominant character in the context of this simple bacterial community (Figure 4B). Interestingly, the IgA^low^ *B. ovatus* strain Q also dominated the relative abundance of the microbiota, as it represented ~95% of the microbiota compared with ~5% of *B. ovatus* strain E (Figure 4C). In an attempt to overcome this resistance to transfer of the high-IgA phenotype to mice with low-IgA phenotype, we performed a similar experiment but added three more IgA^high^ *B. ovatus* strains. Under these conditions, the high-IgA phenotype was transfered to the cohoused mice initially monocolonized with the IgA^low^ strain (Figure S8A). However, *B. ovatus* strain Q still represented a substantial proportion (32.5 - 53.8%) of the relative abundance in this bacterial community (Figure S8B). Thus, a multiplex cocktail of bacterial effector strains that each individually can induce a specific phenotype provides a more robust strategy for transferring a high-IgA phenotype.

**Figure 4.**
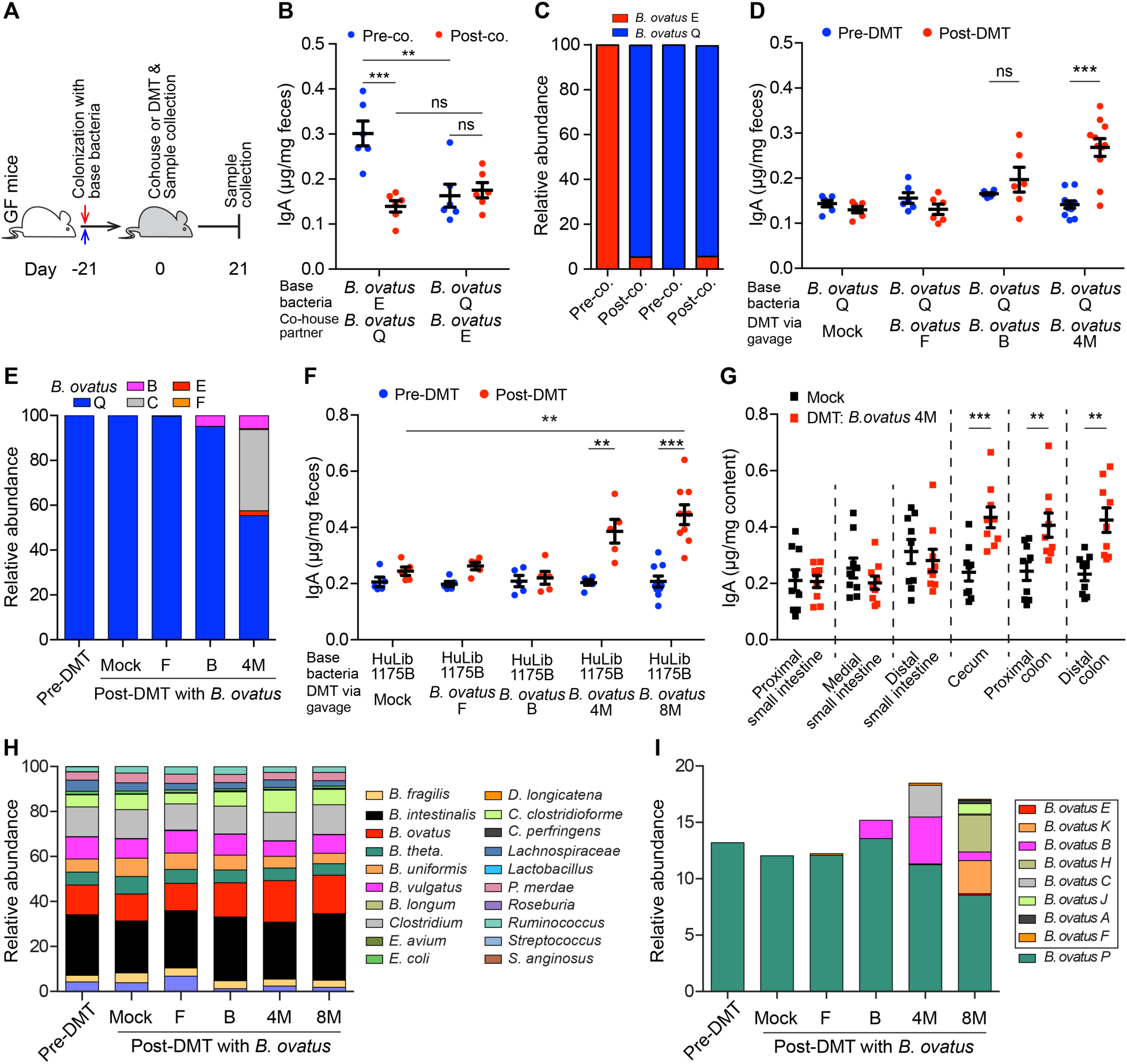
Multiplex microbial strains robustly transfer high-IgA phenotype to low-IgA producing mice. (**A**) Schematic representation of cohousing and defined microbial transplant (DMT) strategies. (**B** and **C**) Fecal IgA concentration (**D**) and relative abundance of each *B. ovatus* strain (**E**) in pre- and post-cohoused gnotobiotic mice, which were pre-colonized with either *B. ovatus* strain E or Q. (**D** and **E**) Fecal IgA concentration (**D**) and relative abundance of each *B. ovatus* strain (**E**) in mice pre- and post-microbial transplantation. Mice were first colonized with *B. ovatus* strain Q for three weeks and subsequently administered cultured microbes comprised of either an individual IgA^high^ *B. ovatus* strain or a cocktail of IgA^high^ *B. ovatus* strains. (**F**) Fecal IgA concentration in mice pre- and post-DMT, which were pre-colonized with human microbiota arrayed culture collection (i.e. HuLib1175B) for three weeks. The administered microbes were either an individual IgA^high^ *B. ovatus* strain or a cocktail of IgA^high^ *B. ovatus* strains. Mock: PBS; *B. ovatus* 4M: a cocktail of 4 different IgA^high^ *B. ovatus* strains; *B. ovatus* 8M: a cocktail of 8 different IgA^high^ *B. ovatus* strains. (**G**) Free IgA concentration along the intestinal tract of mice after DMT with Mock or *B. ovatus* 4M. (**H**) Relative abundance of bacterial species in mice pre- and post-DMT. (**I**) Relative abundance of different *B. ovatus* strains in mice pre- and post-DMT. Data shown are mean ± standard error of the mean. Sequencing plots display the average relative abundance of bacteria from five mice. Each dot represents a biological replicate. Detailed strain information is listed in Tables S2 and S6. *p*-values with statistical significance (assessed by two-tailed Student’s *t* test) are indicated: \**p* < 0.05, \*\**p* < 0.01, \*\*\**p* < 0.001; ns, not significant.

Beyond cohousing, we further validated the above findings by transferring IgA^high^ strains by oral gavage. Consistent with the cohousing results, mice first colonized with an IgA^low^ *B. ovatus* strain and then given a defined microbial transplant (DMT) via oral gavage with an additional IgA^high^ strain did not alter gut IgA secretion. In contrast, mice receiving a cocktail of four IgA^high^ *B. ovatus* strains (*B. ovatus* 4M) produced significantly more fecal IgA (Figure 4D and Table S4). Metagenomic sequencing results demonstrated that multiple *B. ovatus* strains colonized the recipient mice (Figure 4E). Of note, IgA^low^ *B. ovatus* strain Q still dominated the relative abundance of the gut microbiota in individual strain transfers (Figure 4E). IgA^high^ strains accounted for 44% of the gut microbiota in the *B. ovatus* 4M DMT group with each individual IgA^high^ strain having a distinct relative abundance (Figure 4E). Finally, we replicated these results in mice pre-colonized with another IgA^low^ *B. ovatus* strain *R* (Figures S8C and S8D).

To validate these results in the setting of more complex gut microbiotas, we performed similar experiments using either gnotobiotic mice pre-colonized by a synthetic cocktail of diverse bacterial species that included *B. ovatus* IgA^low^ strain Q (Table S5) or gnotobiotic mice pre-colonized with arrayed culture collections established from donors harboring a functionally validated IgA^low^ *B. ovatus* strain (Figure 1H and Table S6). As with simpler communities, transfer of the high-IgA phenotype was robustly achieved with *B. ovatus* 4M or a multiplex cocktail of eight IgA^high^ *B. ovatus* strains (*B. ovatus* 8M) (Table S4) but not by individual IgA^high^ *B. ovatus* strains (Figures 4F and S8E). Consistent with our previous findings, IgA was elevated only in the large intestine (Figures 4G and S8F). The relative proportions of each IgA^high^ strain and total relative abundance of all IgA^high^ strains in the stool of multiplex bacterial cocktail recipient mice varied across recipient microbiota communities (Figures 4H, 4I, S8G and S8H).

Since *B. ovatus* strain C had a high relative abundance after DMT with *B. ovatus* 4M across multiple recipient microbiotas, we examined whether this strain itself could convert low-IgA producing mice to high-IgA producing mice. In receipt mice pre-colonized with either *B. ovatus* strain Q or microbiota arrayed culture collection (i.e. HuLib1175B), transplantation of *B. ovatus* strain C alone did not significantly increase fecal IgA on its own (Fig. 4d,f and Supplementary Fig. 9a,b).

To further validate the IgA-inducing properties of our multiplex IgA^high^ *B. ovatus* cocktails, we tested these cocktails in two additional gnotobiotic mouse models pre-colonized by human microbiota arrayed culture collections with low-IgA potential (Table S6). Again, we found that the multiplex IgA^high^ *B. ovatus* cocktails robustly increased fecal IgA (Figures S10A-S10F). Across all of the tested *B. ovatus* 4M and *B. ovatus* 8M recipients, we did not find a correlation between the total relative abundance of IgA^high^ strains and the fecal IgA levels, which indicates that maximizing the total abundance of IgA^high^ *B. ovatus* strains does not necessarily increase gut IgA production (Figure S10G-S10I). In summary, our results demonstrate that transfer of multiplex IgA^high^ *B. ovatus* strain cocktails, but not that of individual IgA^high^ strains, consistently and robustly modulates the immune system (e.g. IgA phenotype) across several complex pre-existing gut microbiota.

## Discussion

Functional differences of pathogenic bacteria at the strain level have been intensively studied in the past decades and are a fundamental component of infectious disease clinical practice. More recently the functional impact of bacterial strain variation is becoming apparent in the context of the protective or disease-enhancing properties of the commensal microbiota (Arthur et al., 2012; Britton et al., 2019; Palm et al., 2014; Palmela et al., 2018; Viladomiu et al., 2017). Here, we identified that approximately half of the isolated strains from *B. ovatus* species, which is one of the most common species of our gut commensal microbiota, drive increased IgA production in the distal intestinal tract. Interestingly, we did not find that the variation in fecal IgA induced by different *B. ovatus* strains was related to unique genetic lineages amongst strains or the density of the bacteria in the feces. Through manipulation of the pre-existing gut microbiota composition, we discovered that cocktails of IgA^high^ *B. ovatus* strains were more efficient than individual IgA^high^ *B. ovatus* strains in converting mice with low gut IgA production into mice producing large amounts of gut IgA.

IgA^high^ *B. ovatus* strains increased IgA production in distal but not proximal intestinal segments by enhancing the ratio of IgA-secreting B cells. Remarkably, this induction was not dominated by the migration of IgA^+^ B cells from canonical IgA inductive sites, as gnotobiotic mice colonized with either IgA^high^ or IgA^low^ *B. ovatus* strains showed comparable IgA^+^ B cells in PPs and MLNs. One possibility is that IgA^high^ *B. ovatus* strains locally elicit IgA production in the large intestine including cecal patches, ILFs and LP (Cerutti and Rescigno, 2008; Fagarasan et al., 2010; Masahata et al., 2014). Interestingly, mice harboring specific *B. ovatus* strains showed no significant differences in the intestinal abundance of *B. ovatus*, which further highlights the unique IgA-inducing properties of individual strains.

After IgA^high^ *B. ovatus* strain colonization, CD4^+^ T cell-depleted mice showed a reduced ratio of IgA^+^ B cells in the gut, in turn leading to decreased luminal IgA significantly along the entire intestinal tract. Of note, both CD4^+^ T cell-sufficient and T cell-depleted mice produced a comparable level of gut IgA at the beginning post colonization, which suggests CD4^+^ T cells play less of a role during the very early stage of IgA induction likely due to the dominance of the TI B cell-activation pathway. Interestingly, a protein from the gut commensal *Lactobacillus rhamnosus* was recently shown to locally elicit IgA production via gut epithelial cells (Wang et al., 2017). Thus, further studies will be needed to delineate the precise mechanisms whereby IgA^high^ *B. ovatus* strain colonized mice generate gut IgA. Nevertheless, our study highlights the important contribution of CD4^+^ T cells in bacteria-mediated IgA production, especially in the large intestine (Kawamoto et al., 2014; Kunisawa et al., 2013).

FMT has a high success rate in the treatment of recurrent *C. difficile* infection (van Nood et al., 2013). However, its success in other indications, such as ulcerative colitis, is more limited (Moayyedi et al., 2015; Rossen et al., 2015). Although improving bacteria engraftment remains a key goal of microbiota manipulation (Grinspan and Kelly, 2015; Moayyedi et al., 2015; Rossen et al., 2015), identifying new strategies that optimize the transfer of a specific immune phenotype constitutes a goal with a potentially larger range of applications. Using IgA induction as an example of immunomodulatory phenotype transfer, our data showed that multiplex bacterial cocktails of IgA^high^ *B. ovatus* strains elicited a more robust phenotype transfer than any individual strain, even in mice with complex gut ecosystem. This multiplex effector strain cocktail strategy was robust across multiple recipients, pre-colonized with different low-IgA microbiotas, and could represent an effective approach to modify gut immune parameters in addition to IgA. Of note, across the tested inductions of IgA via gut microbiota manipulation with IgA^high^ *B. ovatus* strains, we found that no single strain consistently dominated over the others. Thus, multiplex bacterial cocktails do not appear to have “super strains” with dominant IgA-inducing function. Rather, the combination of multiple IgA^high^ effector strains in these cocktails has an IgA-inducing potential superior to that of any individual strain. Intriguingly, the relative abundance of total *B. ovatus* species remained largely stable even after the introduction of one to eight new strains suggesting that these new strains largely share the same ecological niches as that occupied by the pre-existing *B. ovatus*.

Gut microbiota-based immunomodulation has shown great potential as a therapeutic in a number of mouse models, including Treg induction to limit colitis (Atarashi et al., 2013), mitigation of graft-versus-host disease (Mathewson et al., 2016), and alteration of immune checkpoint inhibitor efficacy for immuno-oncology (Tanoue et al., 2019). As immunotherapeutic bacterial cocktails move towards the clinic (Honda and Littman, 2016), a key factor in their success will be robust and consistent manipulation of the desired immune populations. The usage of multiplex cocktails with each strain independently capable of inducing the desired immune modulation provides one potential route to a consistent immune response that is robust to the variation in microbiome composition across individuals.

In summary, our results highlight the importance of bacterial strain variation on the IgA-inducing potential of the gut microbiota. In addition, we also identify a new strategy (i.e. multiplex bacterial strain cocktail) for the exploitation of strain variation in the development of robust microbiota-based immunomodulation strategies.

## Methods

### Mice

Germ-free C57BL/6 and Swiss Webster mice were bred and maintained in flexible film gnotobiotic isolators (Class Biologically Clean, Ltd.). All mice were group housed with a 12-hour light/dark cycle and allowed *ad libitum* access to diet and water. All animal studies were carried out in accordance with protocols approved by the Institutional Animal Care and Use Committee (IACUC) in Icahn School of Medicine at Mount Sinai.

### Colonization of germ-free mice with cultured bacteria

Germ-free mice (~8 weeks old) were colonized 200-μl aliquot of bacteria suspension via oral gavage. Colonized mice were housed in flexible film vinyl isolators or in filter top cages using previously described techniques (Faith et al., 2013).

### Growth and isolation of bacterial strains

All bacterial strains were obtained from previously banked stool, public culture repositories or human gut microbiota arrayed culture collections (Faith et al., 2014). All bacterial strains isolated for this study were isolated from deidentified stool samples from individuals under a Mount Sinai IRB approved protocol (IRB-16-00008). All bacteria apart from *E. coli* were grown under anaerobic condition at 37°C in Brain Heart Infusion medium supplemented with 0.5% yeast extract (Difco Laboratories), 0.4% monosaccharide mixture, 0.3% disaccharide mixture, L-cysteine (0.5 mg/ml; Sigma-Aldrich), malic acid (1 mg/ml; Sigma-Aldrich) and 5 μg/ml hemin. *E. coli* was cultured in LB Broth Miller (EMD Chemicals, Inc.) under aerobic condition at 37°C.

### Quantification of immunoglobulin A by ELISA

Total fecal IgA were measured by sandwich ELISA. High-binding ELISA plates (Corning 3690) were coated with 1 μg/ml goat anti-mouse IgA (SouthernBiotech, AL) capture antibody overnight at 4°C. Plates were washed and blocked with 1% BSA in PBS for 2 h at room temperature. Diluted samples and standards were added and incubated overnight at 4°C. Captured IgA was detected by horseradish peroxidase (HRP)-conjugated goat anti-mouse IgA antibody (Sigma-Aldrich). ELISA plates were developed by TMB microwell peroxidase substrate (KPL, Inc.) and quenched by 1 M H_2_SO_4_. Colorimetric reaction was measured at OD = 450 nm by a Synergy™ HTX Multi-Mode Microplate Reader (BioTek Instruments, Inc.). For the quantification of human stool IgA, the same ELISA procedure as described above was performed except using anti-human IgA and anti-human IgA-HRP antibodies (SouthernBiotech, AL). Corresponding immunoglobulin isotypes were used as standards after serial dilutions.

### Detection of IgA-coated bacteria in feces

IgA-coated fecal bacteria were measured by flow cytometry as previously described (Kau et al., 2015; Palm et al., 2014). Briefly, mouse fecal pellets, stored at −80°C freezer after collection, were dissolved in PBS to a final concentration of 100 milligram per milliliter PBS by weight, thawed at room temperature, homogenized in vortex mixer and centrifuged at 4°C to remove large particles. The supernatant was passed through a 40 μm sterile nylon filter and a small aliquot of the bacteria suspension was collected for staining. Bacteria were pelleted by centrifugation and washed in PBS/1%BSA/2mM EDTA buffer for 3 times. Non-specific binding sites were first blocked with 50 μl 20% rat serum for 20 min at 4°C. Bacteria were then stained with monoclonal rat anti-mouse IgA antibody (eBioscience, clone mA-6E1) for 30 min at 4°C. After washing 3 times, bacterial pellets were resuspended in PBS containing SYBR Green I (Invitrogen, USA). Samples were run through a BD LSR Fortessa™ cell analyzer and further analyzed by FlowJo software (Tree Star, Inc.). Only SYBR positive events were regarded as real bacteria and gated for further quantification of IgA-coated bacteria (Figure S11A).

### Lymphocyte isolation from tissues

To isolate mononuclear cells from Peyer’s patches (PPs), PPs were excised from mouse small intestines and incubated in dissociation buffer, containing Hank’s Balanced Salt Solution (HBSS) without Ca^2+^ and Mg^2+^ (GIBCO), 10% fetal bovine serum (FBS), 5 mM EDTA and 15 mM HEPES, at 37°C for 30 min. Later, tissues were mechanically separated by pushing them through a 70 μm strainer into Iscove’s Modified Dulbecco’s Medium (IMDM) supplemented with 2% FBS. Filtered cells were spun down, washed and resuspended in IMDM/2%FBS. Lamina propria lymphocytes were isolated as described (Faith et al., 2014). Briefly, small intestines and colons were excised, followed by removing visceral fat and intestinal contents. Tissues were opened longitudinally, washed twice in HBSS and incubated in dissociation buffer for 30 min at 37°C with mild agitation to remove epithelium and intraepithelial lymphocytes. Tissues were then washed three times in ice cold HBSS, cut into ~2 cm pieces and digested with collagenase (Sigma-Aldrich), DNase I (Sigma-Aldrich) and dispase I (Corning). Cell suspensions were filtered through 70 μm cell strainers, washed three times, and resuspended in IMDM/2%FBS. Mesenteric lymph nodes were separated from mesenteric fat and dissociated in IMDM/2%FBS by physically pressing the tissues between the frosted portions of two glass microscope slides. The cell suspension was filtered through a 70 μm cell strainer, washed three times and resuspended in IMDM/2%FBS.

### Flow cytometry analysis and antibodies

Isolated mononuclear cells were washed in PBS and incubated with Zombie Aqua™ dye (BioLegend) to distinguish live and dead cells. Before surface staining, non-specific binding of immunoglobulin to Fc receptors was blocked by anti-mouse CD16/32 antibody (BD Biosciences). Cells were stained in FACS buffer (PBS without Ca^2+^/Mg^2+^ supplemented with 2% FBS and 2 mM EDTA) containing a mix of antibodies for 30 min at 4°C. The following antibodies were purchased from BioLegend if not indicated otherwise: anti-mouse CD45 (clone 30-F11), anti-mouse/human CD45R/B220 (clone RA3-6B2), anti-mouse GL7 (clone GL7), anti-mouse CD4 (clones GK1.5 and RM4-4), anti-mouse IgA (eBioscience, clone mA-6E1). For the staining of IgA^+^ cells, both surface and intracellular staining were performed. Multi-parameter analysis was conducted with BD^TM^ LSR II flow cytometry or BD LSR Fortessa™ cell analyzers and analyzed with FlowJo software (Tree Star, Inc.). Only singlets and live cells were used in all further analyses (Figure S11B).

### Immunofluorescence staining

Immunofluorescence staining was performed as described previously (Moon et al., 2015). Briefly, intestinal tissues were fixed in 10% neutral formalin overnight at 4°C, dehydrated in 15% and 30% sucrose buffer sequentially and mounted in O.C.T Embedding Compound (Electron Microscopy Sciences). Cryostat sections (~8 μm) were prepared, blocked with anti-CD16/32 antibody in 10% (v/v) rat serum/0.1% Triton-X100 in PBS for 30 min at room temperature and incubated with the indicated primary antibodies at 4°C overnight. The following primary antibodies were used: rat anti-mouse IgA-FITC (eBioscience, clone mA-6E1), goat anti-mouse pIgR (R&D Systems, cat #: AF2800). Slides were washed in PBS for three times, incubated with Alexa Fluor^®^-conjugated species-specific secondary antibody (Invitrogen) for 1 h at room temperature if needed and finally mounted with ProLong^®^ Gold Anti-fade Reagent with DAPI (Invitrogen). Fluorescence images of sections were acquired with a LSM780 confocal laser-scanning microscope (Carl Zeiss) and further processed in ImageJ if necessary.

### Depletion of CD4^+^ T cells in germ-free mice

*In vivo* depletion of CD4^+^ T cells was performed as described (Kruisbeek, 2001). Briefly, germ-free mice (8 weeks old) were first injected intraperitoneally (i.p.) with anti-mouse CD4 monoclonal antibody (Bio X Cell, clone GK1.5) or matched isotype control (Bio X Cell, clone LTF-2) at 0.5 mg/day/mouse for 3 consecutive days. Five days after the first antibody injection, mice were inoculated via oral gavage with *B. ovatus* strain E or Q. Then the injection was performed every 3 days for a period of 3 weeks. Efficacy of T cell depletion in gnotobiotic mice before and after *B. ovatus* colonization was evaluated by flow cytometry.

### Extraction of bacterial DNA from feces

Each murine fecal pellet was collected into a 2 ml screw cap tube (Axygen Scientific, SCT200SSC) and stored at −80°C freezer until processing. Each sample was mixed with 1.3 ml of buffer, composed of 282 μl of DNA buffer A (20 mM Tris pH 8.0, 2 mM EDTA and 200 mM NaCl), 200 μl of 20% SDS (v/w), 550 μl of Phenol:Chloroform:IAA (25:24:1) (Ambion, AM9732) and 268 μl of Buffer PM (Qiagen, 19083), and 400 μl of 0.1 mm diameter zirconia/silica beads (BioSpec, 11079101z). Next, the sample was mechanically lysed with a Mini-Beadbeater-96 (BioSpec, 1001) for 5 min at room temperature. After centrifuging for 5 min at 4000 rpm (Eppendorf Centrifuge 5810 R), all aqueous phase was collected, mixed with 650 μl of Buffer PM thoroughly before running through a Qiagen spin column. The column was washed twice with Buffer PE (Qiagen, 19065). Attached DNA was eluted with 100 μl of Buffer EB (Qiagen, 19086) and quantified with Qubit^TM^ dsDNA Assay Kit (Thermo Fisher Scientific, Q32853/Q32854). Bacteria density was calculated by the following equation: Bacteria Density = DNA yield per sample (ug) / weight of sample (mg) (Contijoch et al., 2019).

### Bacterial genome and metagenomic sequencing

Purified bacterial template DNA (~250 ng) was sonicated and prepared using the NEBNext^®^ Ultra^™^ II DNA Library Prep kit. Samples were pooled and sequenced with an Illumina HiSeq 4000 with pair-end 150nt reads. Metagenomic sequencing reads were mapped back to the reference genomes for each experiment to determine the relative abundance of each strain. To uniquely distinguish each strain, 100K sequencing reads for each sample were mapped to the unique regions of each genome and final abundances were scaled by the unique genome size of each strain (i.e. genome equivalents), as previously described (McNulty et al., 2013).

### Statistical analysis

Data are shown as mean ± SEM. Statistical significance between two groups was assessed by an unpaired, two-tailed Student’s *t* test. Comparisons among three or more groups were performed using One-way ANOVA. Bimodality distribution of IgA levels induced by different *B. ovatus* strains was performed in R (R package ‘diptest’). For correlation test, Pearson correlation coefficient was employed. Data plotting, interpolation and statistical analysis were performed using GraphPad Prism 6.0 (GraphPad Software, La Jolla, CA) or R statistical software (version 3.2.2). A *p*-value less than 0.05 is considered statistically significant.

### Data and code availability

Bacterial genomes and metagenomic sequencing reads for this study are available via NCBI BioProject accession number PRJNA518912.

## Acknowledgments

We are grateful to C. Fermin, E. Vazquez, and G. Escano for the husbandry of gnotobiotic mice; Drs. Anuk A. Das, Dirk D. Gevers, Charlotte Cunningham-Rundles, Brian Brown and Thomas Moran for helpful discussions and comments. This work was supported in part by the staff and resources of the Gnotobiotic Mouse Core Facility, the Microbiome Translational Center, the Flow Cytometry Core Facility, the Microscopy CoRE and the Scientific Computing Division in Icahn School of Medicine at Mount Sinai. This work was supported by National Institutes of Health Grants (NIGMS GM108505, NCCIH AT008661, NIDDK DK112978) and Janssen Human Microbiome Institute (to J.J.F.) and NIH DK112679 (to E.J.C.).

## Author contributions

C.Y. and J.J.F conceived the study and designed the experiments; C.Y., I.M., E.J.C., J.B., V.A., S.S., E.G., D.H., M.D. and J.J.F. collected samples and conducted the experiments; I.M. and Z.L. provided bacterial isolates; C.Y., S.M., A.C. and J.J.F. analyzed data; C.Y. and J.J.F. prepared the manuscript. All authors read and approved the final manuscript.

