## Supplemental Information for "Strain-level differences in gut microbiome composition determine fecal IgA levels and are modifiable by gut microbiota manipulation"

#### Supplemental methods

**Quantification of immunoglobulins by ELISA**

Like the quantification of total fecal IgA, serum immunoglobulin isotypes (IgA, IgG1, IgG2a, IgG2b, IgG3, IgM and IgE) were also detected using sandwich ELISA following the same procedures as the quantification of fecal IgA except for using different capture and detection antibody pairs (all following antibodies were purchased from Southern Biotechnology Associates, Inc. if not indicated otherwise): goat anti-mouse IgA, goat anti-mouse IgG1, goat anti-mouse IgG2a, goat anti-mouse IgG2b, goat anti-mouse IgG3, rat anti-mouse IgE, rat anti-mouse IgM, goat anti-mouse IgG-HRP, goat anti-mouse IgE-HRP, goat anti-mouse IgM-HRP and goat anti-mouse IgA-HRP (Sigma-Aldrich). Corresponding mouse immunoglobulin isotypes were used as standards after serial dilutions.

**RNA isolation**

Excised small intestine and colon from gnotobiotic mice that were colonized for three weeks with *B. ovatus* E or Q strains were kept in RNA*later* RNA Stabilization Reagent (Qiagen, 76104) and stored at -20°C freezer until future processing. Total RNA was extracted with the RNeasy Mini Kit (Qiagen, 74104) according to manufacture's protocol. The concentration and quality of RNA was analyzed with a NanoDrop™ 8000 Spectrophotometer (Thermo Fisher Scientific, USA).

**Quantification of mRNA by quantitative RT-PCR**

The cDNA for each sample was synthesized with High-Capacity cDNA Reverse Transcription Kits (Applied Biosystems, 4368813). The StepOne Real-Time PCR System (Applied Biosystems, USA) was used for PCR amplification of the cDNAs with Applied Biosystems™ Power SYBR™ Green Master Mix and oligonucleotide primer pairs specific for pIgR, Muc2 and glyceraldehydes-3-phosphate dehydrogenase (GAPDH) mRNAs. The primer sequences were as follows: GAPDH forward primer, 5'-TGAACGGGAAGCTCACTGG-3'; GAPDH reverse primer, 5'-TCCACCACCCTGTTGCTGTA-3'; pIgR forward primer, 5'-AGGCAATGACAACATGGGG-3'; pIgR reverse primer, 5'-ATGTCAGCTTCCTCCTTGG-3' (Nakamura et al., 2012); Muc2 forward primer, 5'-GCTGACGAGTGGTTGGTGAATG-3'; Muc2 reverse primer, 5'-GATGAGGTGGCAGACAGGAGAC-3' (Wlodarska et al., 2011). The following parameters were set for cDNA amplification and quantification: 30 seconds at 95°C, and then 40 cycles of denaturation at 95°C for 15 seconds and annealing at 60°C for 1 minutes. The mRNA level of test gene was normalized to GAPDH according to the formula: $(2^{-(C_T^{test} - C_T^{GAPDH})}) \times 100\%$.

**Scanning electron microscopy**

The morphology of *B. ovatus* in mouse colonic tissue was observed under scanning electron microscopy (SEM). Colon tissues were excised from gnotobiotic mice colonized for three weeks with either *B. ovatus* strain E or Q and fixed in 3% glutaraldehyde buffer overnight at 4°C. Samples were then washed gently in 0.2 M sodium cacodylate buffer to remove residual fixative and re-fixed with 1% osmium tetraoxide/0.2 M cacodylate buffer for one hour. After complete drying, samples were first coated with gold particles and observed with a HITACHI S-4300 SEM (HITACHI, Japan).

**Treatment of gnotobiotic mice with FTY720**

Germ-free mice were administered 2-Amino-2-[2-(4-octyl-phenyl)-ethyl]-propane-1,3-diol hydrochloride (FTY720) (Sigma-Aldrich, SML0700) by i.p. at 1 µg/g body weight, as previously described (Kunisawa et al., 2007; Ruane et al., 2013). Three days after the initial treatment, mice were colonized with *B. ovatus* strain E and injected FTY720 by i.p. followed by treatment with FTY720 every three days for a period of three weeks. PBS was used in control mice. At the end of the experiment, content from different regions of the intestinal tract was harvested and subject to IgA quantification by ELISA.

### Supplemental results

#### $IgA^{high}$ *B. ovatus* strains induce comparable level of fecal IgA to Taconic SPF microbiota

Segmented filamentous bacterium (SFB) has been described as a potent fecal IgA inducer (Talham et al., 1999). As a reference to compare with IgA levels induced by diverse human gut microbial strains, we measured fecal IgA in C57Bl/6 mice from three mouse vendors. Taconic SPF mice, that are known to be colonized by SFB (Ivanov et al., 2009), produced more IgA in their stool than other mice (i.e. JAX SPF mice and Charles River SPF mice) (Figure S2D). However, the concentration of stool IgA in Taconic mice was comparable to that of gnotobiotic mice colonized with a single $IgA^{high}$ *B. ovatus* strain (Figure 1F and S2D) suggesting that *B. ovatus* $IgA^{high}$ stains are as efficient as SFB in fecal IgA induction in mice.

#### *B. ovatus* induce development of IgA-secreting cells locally in the large intestine

Well-organized follicular structures such as Peyer's patches (PPs) in the small intestine are the most prominent IgA inductive sites. We determined whether $IgA^{high}$ *B. ovatus*-induced IgA-secreting cells residing in the LP of the large intestine had emerged from the small intestinal IgA inductive sites (e.g. PPs). To address this question, we took advantage of FTY720, an S1PR1 agonist, which blocks the cellular egress from secondary lymphoid tissues (Kunisawa et al., 2007). We found that, in general, FTY720-treated and untreated control mice generated comparable fecal IgA after three weeks of colonization (Figure S12A). However, the treated mice, compared to controls, had significantly less luminal IgA in the distal small intestine but not in the large intestinal regions (Figure S12B). These results suggest local development of IgA-secreting B cells in the colon might be the mechanism driving elevated colonic IgA induced by

IgA$^{high}$ strains or that IgA$^{high}$ *B. ovatus* strains promote the survival of IgA-secreting cells in the colon (Fagarasan et al., 2010; Masahata et al., 2014).

### Supplemental figures

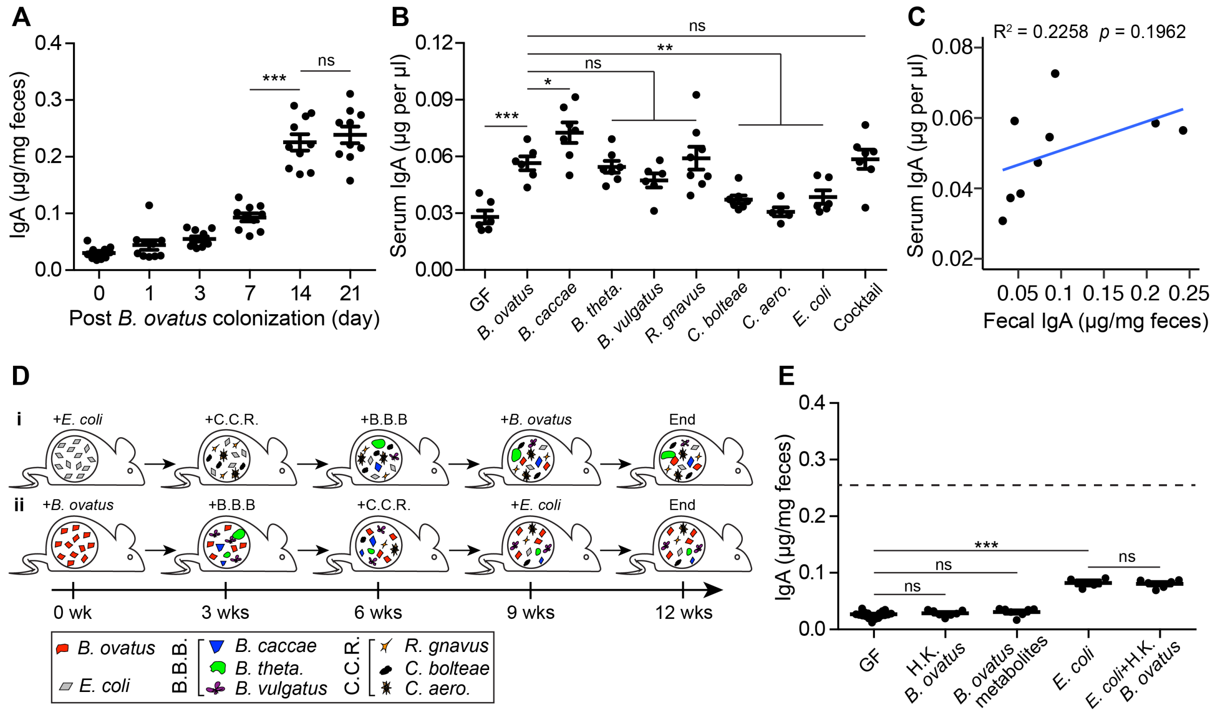


**Figure S1. *B. ovatus* species do not induce more serum IgA production than other bacterial species, related to Figure 1.** (**A**) Fecal IgA dynamics in gnotobiotic mice after colonization with *B. ovatus* strain E. (**B**) Total serum IgA concentration in gnotobiotic mice that were colonized with individual or a cocktail of eight bacterial species for three weeks. (**C**) Correlation of IgA concentration in stool and serum in mice inoculated with different bacterial species. The average concentrations of stool IgA in Figure 1A and serum IgA in Figure S1B were used for plotting. (**D**) Gnotobiotic mice were serially colonized with different bacteria every three weeks. Before each new bacteria addition, stool samples were collected for further analysis. (**E**) Fecal IgA concentration in mice treated with either heat-killed (H.K.) *B. ovatus* or *B. ovatus* metabolites (i.e. filtered, conditioned growth medium from stationary phase of *B. ovatus* cultures). The right side of this plot shows fecal IgA concentration in *E. coli*-precolonized gnotobiotic mice, which were then treated with H.K.-*B. ovatus*. Either metabolites of *B. ovatus* in culture medium or H.K.-*B. ovatus* was used to feed mice, accordingly, for the duration of the experiments. Dotted line indicates the average level of stool IgA induced by viable *B. ovatus*

strain E. Data shown are mean ± standard error of the mean. Each dot represents a biological replicate. Detailed strain information is listed in Table S1. $p$-values with statistical significance (assessed by two-tailed Student's $t$ test) are indicated: ***$p < 0.001$; ns, not significant.

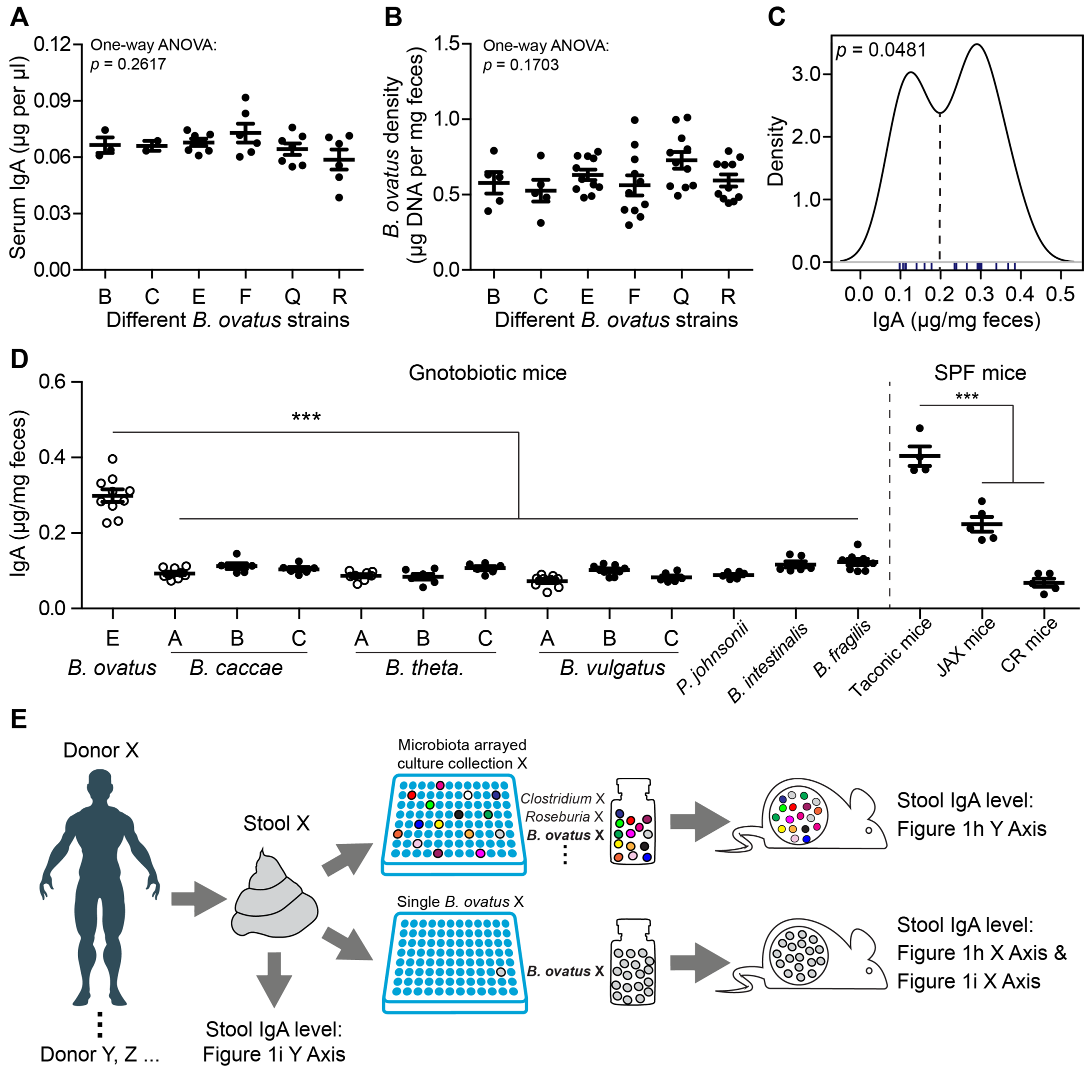


**Figure S2. Strain-level variation in fecal IgA induction was not observed in other tested bacterial species, related to Figure 1.** (**A**) Total serum IgA level in gnotobiotic mice harboring different strains of *B. ovatus*. (**B**) *B. ovatus* density in the stool of gnotobiotic mice colonized with different *B. ovatus* strains. (**C**) Binomial distribution of *B. ovatus* strains in IgA induction. (**D**) Fecal IgA concentration in mice colonized with different strains of *B. caccae*, *B. theta.*, *B. vulgatus* and other Bacteroidales, such as *P. johnsonii*, *B. intestinalis* and *B. fragilis* and in SPF B6 mice purchased from different vendors. Taconic mice were purchased from Taconic Biosciences; JAX mice were purchased from The Jackson Laboratory and CR mice were purchased from Charles River Laboratories. Data with open circle were replotted from Figure 1A to facilitate comparison. (**E**) Experiment schema of the Figures 1H and 1I. Data shown are mean ± standard error of the mean. Each dot represents a biological replicate. Detailed strain information is listed in Tables S2 and S7. $p$-values with statistical significance (assessed by two-tailed Student's $t$ test or one-way ANOVA) are indicated: ***$p < 0.001$; ns, not significant.

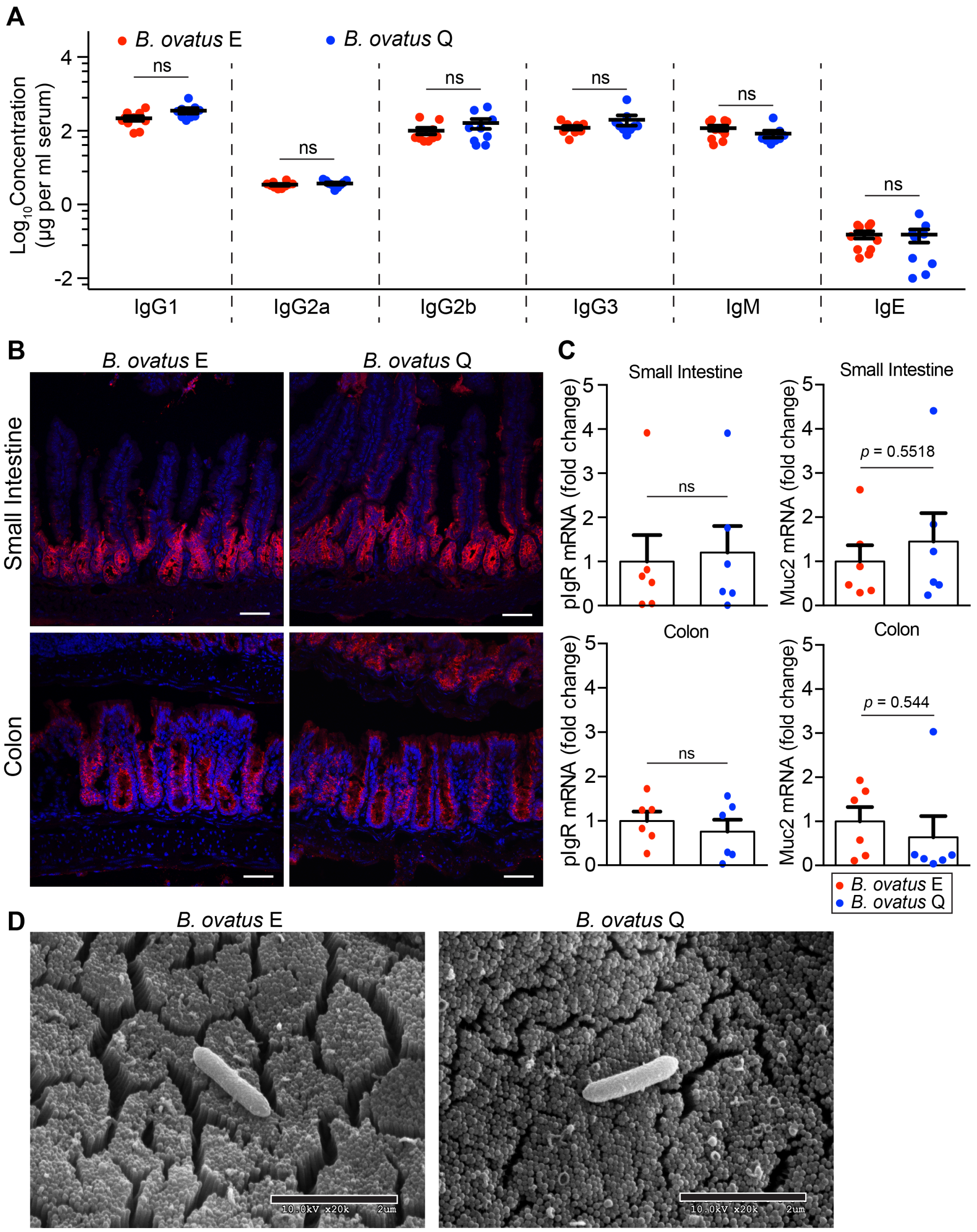


**Figure S3. *B. ovatus* strains E and Q induced comparable levels of different serum immunoglobulin isotypes, pIgR and Muc2 expression in both small intestine and the colon, related to Figure 2.** (**A**) Total serum IgG1, IgG2a, IgG2b, IgG3, IgM and IgE in gnotobiotic mice colonized with either *B. ovatus* strain E or Q. (**B**) Colonic and ileal sections from *B. ovatus* strain E or Q colonized mice were stained with anti-pIgR (Red) and DAPI (4',6-diamidino-2-phenylindole) (Blue). Representative images are shown (n = 4-6 mice per group). Scale Bars = 50 μm. Data shown are mean ± standard error of the mean. (**C**) The fold change of pIgR mRNA level (left) and Muc2 mRNA level (right) in the small intestine and colon of mice colonized for three weeks with *B. ovatus* strain E or Q. (**D**) The morphology of *B. ovatus* strain E or Q in gnotobiotic mice colon. Scale Bars = 2 μm. Each dot represents a biological replicate. *p*-values with statistical significance (assessed by unpaired two-tailed Student's *t* test) are indicated: *$p < 0.05$; ns, not significant.

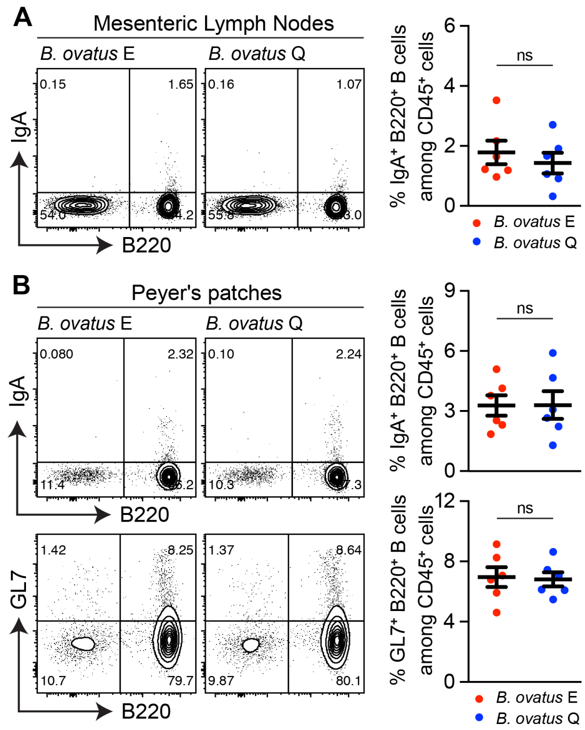


**Figure S4. Quantification of $IgA^{+}B220^{+}$ B cells in MLNs, PPs in *B. ovatus* strain E or Q harboring gnotobiotic mice, related to Figure 2.** (**A**) Representative flow cytometry plot and quantification of $IgA^{+}$ B cells in mesenteric lymph nodes. (**B**) Representative flow cytometry plot and quantification of $IgA^{+}$ B cell and germinal center B cells ($GL7^{+}B220^{+}$) in PPS. Number adjacent to gate represents percentage. Data shown are mean ± standard error of the mean. Each dot represents an individual mouse. *p*-values with statistical significance (assessed by unpaired two-tailed Student's *t* test) are indicated: $^{*}p < 0.05$; ns, not significant.

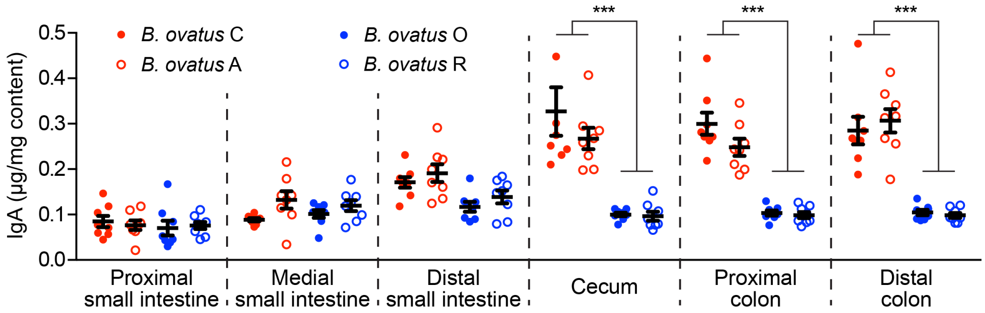


**Figure S5. IgA$^{high}$ *B. ovatus* strains specifically induced more fecal IgA production in the large intestinal regions than IgA$^{low}$ *B. ovatus* strains, related to Figure 2.** Free IgA concentration in different regions along the whole intestinal tract of mice that were colonized with individual *B. ovatus* strains (IgA$^{high}$ *B. ovatus* strains *A* and *C*; IgA$^{low}$ *B. ovatus* strains *O* and *R*) for three weeks. Data shown are mean ± standard error of the mean. Each dot represents a biological replicate. Detailed strain information is listed in Table S2. *p*-values with statistical significance (assessed by unpaired two-tailed Student's *t* test) are indicated: ***$p$ < 0.001; ns, not significant.

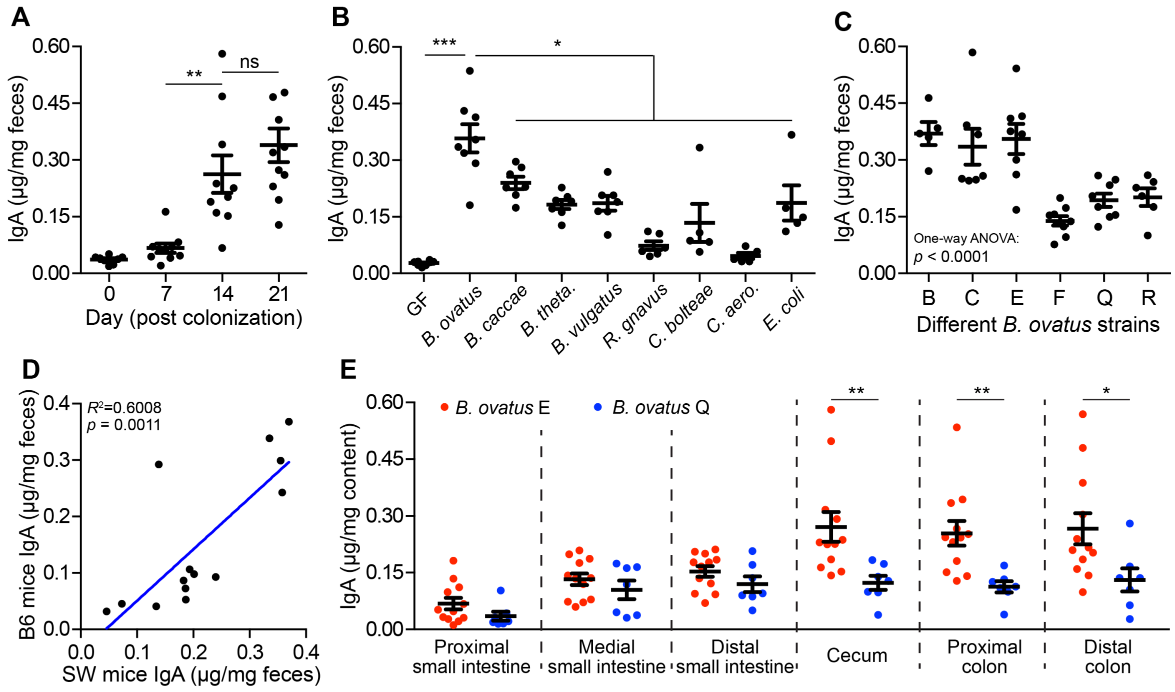


**Figure S6. Host genetic background has little influence on human gut bacteria induced fecal IgA production in mice.** (**A**) Dynamics of fecal IgA concentration in gnotobiotic Swiss Webster mice colonized with *B. ovatus* strain E. (**B** and **C**) Fecal IgA concentration in gnotobiotic Swiss Webster mice colonized with different bacterial species (**B**) or various strains of *B. ovatus* (**C**). (**D**) Correlation of fecal IgA produced by B6 mice and Swiss Webster mice after the same bacteria colonization. The average fecal IgA level of Swiss Webster mice in (**B**) and (**C**) were plotted against the average fecal IgA level in C57BL6/J mice that were colonized with the same bacterial species or strain from Figure 1A and Figure 1F. (**E**) Luminal IgA concentration along the whole intestine in Swiss Webster mice colonized with either *B. ovatus* strains E or Q. Data shown are mean ± standard error of the mean. Each dot represents a biological replicate in **A**, **B**, **C** and **E**. Detailed strain information is listed in Tables S1 and S2. *p*-values with statistical significance (assessed by two-tailed Student's *t* test or one-way ANOVA) are indicated: *$p < 0.05$, **$p < 0.01$, ***$p < 0.001$; ns, not significant.

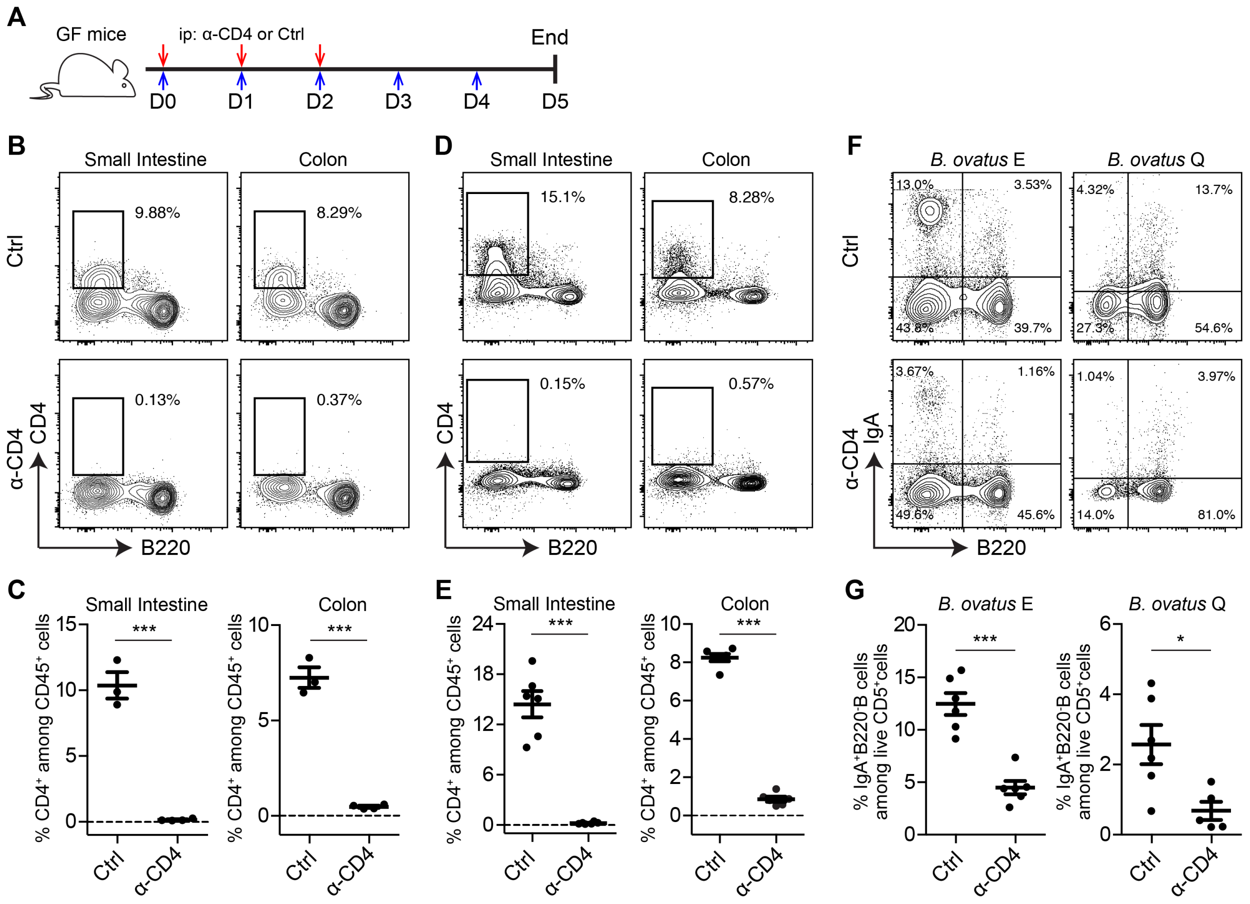


**Figure 7. Anti-CD4 antibody promptly and robustly depleted CD4$^+$ T cells in multiple organs of germ-free and gnotobiotic mice, related to Figure 3.** (**A**) Schematic representation of anti-CD4 antibody injection. GF mice were injected intraperitoneally with anti-CD4 antibody or isotype control for three consecutive days (0.5 mg/mouse/day). Three days after the last injection, tissues were collected and processed. Red arrows indicate antibody injection and blue arrows represent time. (**B** and **C**) Representative flow cytometry plot (**B**) and quantification (**C**) of CD4$^+$ T cells in the LP of small intestine and colon in germ-free mice. (**D** and **E)** Representative flow cytometry plot (**D**) and quantification (**E**) of CD4$^+$ T cells in LP of small intestine and colon of gnotobiotic mice colonized with *B. ovatus* strain E with or without anti-CD4 antibody treatment. (**F** and **G)** Representative flow cytometry plot (**F**) and quantification (**G**) of IgA$^+$B220$^-$ cells in the LP of small intestine of gnotobiotic mice colonized with *B. ovatus* strain E or Q w/o anti-CD4 antibody treatment. Number adjacent to gate

represents percentage. Data shown are mean ± standard error of the mean. Each dot represents a biological replicate. *p*-values with statistical significance (assessed by unpaired two-tailed Student's *t* test) are indicated: *$p < 0.05$, ***$p < 0.001$; ns, not significant.

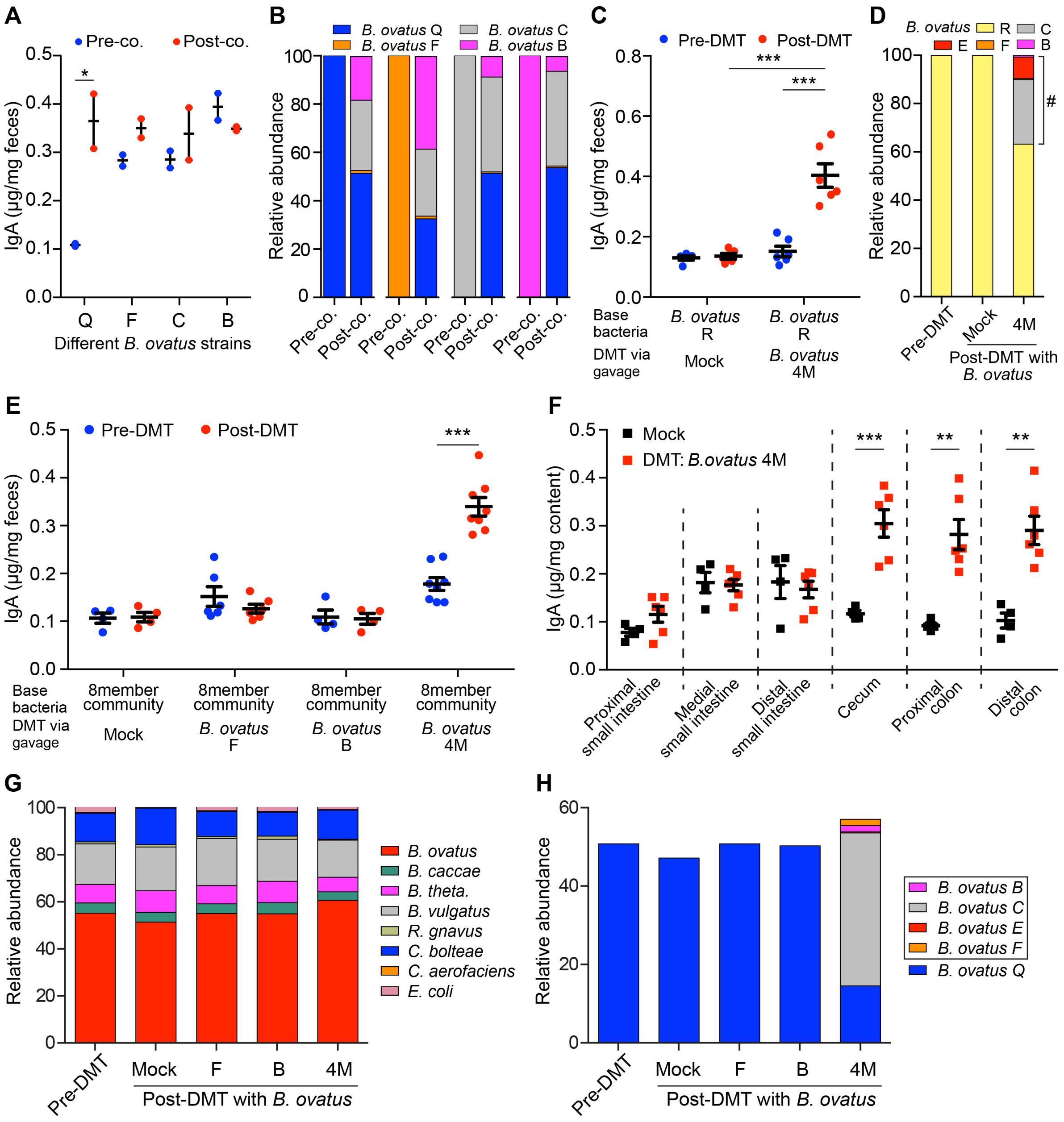


**Figure S8. Multiplex cocktail of microbial strains overcome phenotype transfer resistance in gnotobiotic mice that were pre-colonized with simple bacterial community, related to Figure 4.** (**A** and **B**) Fecal IgA concentration (**A**) and relative abundance of each *B. ovatus* strain (**B**) in pre- and post-cohoused gnotobiotic mice. Before cohousing, four groups of mice were pre-colonized with four individual *B. ovatus* strains, respectively, for three weeks. Then, all mice were cohoused together at a ratio of 1:1:1:1 for another three weeks. (**C** and **D**) Fecal IgA concentration (**C**) and relative abundance of each *B. ovatus* strain (**D**) in mice pre- and post-

DMT. Mice were first colonized with *B. ovatus* strain *R* for three weeks. Then the microbial cocktail *B. ovatus* 4M was administered. (**E**) Fecal IgA concentration in mice pre- and post-DMT, which were pre-colonized with eight-member bacterial community for three weeks. The microbial cocktail consisted of either an individual $IgA^{high}$ *B. ovatus* strain or *B. ovatus* 4M. (**F**) Luminal IgA concentration along the intestinal tract of mice after gavage with Mock (PBS) or *B. ovatus* 4M. (**G**) Relative abundance of each bacterial species in mice pre- and post-DMT. (**H**) Relative abundance of different *B. ovatus* strains in mice pre- and post-DMT. Data shown are mean ± standard error of the mean. Sequencing plots display the average abundance from five mice. Each dot represents a biological replicate. Detailed bacteria information is listed in Tables S2, S4 and S5. *p*-values with statistical significance (assessed by two-tailed Student's *t* test) are indicated: *$p < 0.05$, **$p < 0.01$, ***$p < 0.001$; ns, not significant.

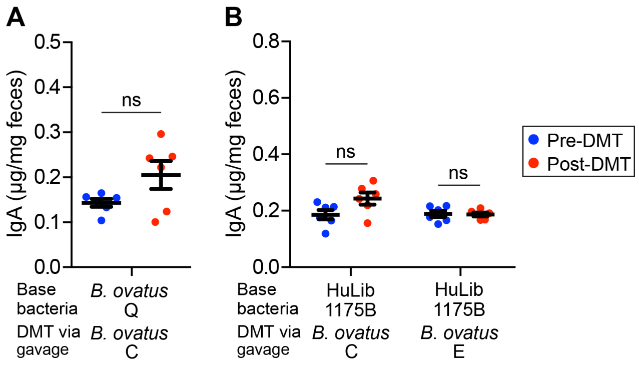


**Figure S9. *B. ovatus* strain C and E individually do not convert low-IgA to high-IgA producing mice, related to Figure 4.** (**A**) Fecal IgA concentration in gnotobiotic mice pre-DMT and post-DMT. Before DMT, mice were pre-colonized with *B. ovatus* strain Q for three weeks. (**B**) Fecal IgA concentration in gnotobiotic mice pre-DMT and post-DMT. Mice were pre-colonized with microbiota arrayed culture collections (i.e. HuLib1175B) for three weeks. Data shown are mean ± standard error of the mean. Each dot represents a biological replicate. *p*-values with statistical significance (assessed by unpaired two-tailed Student's *t* test) are indicated: *$p < 0.05$; ns, not significant.

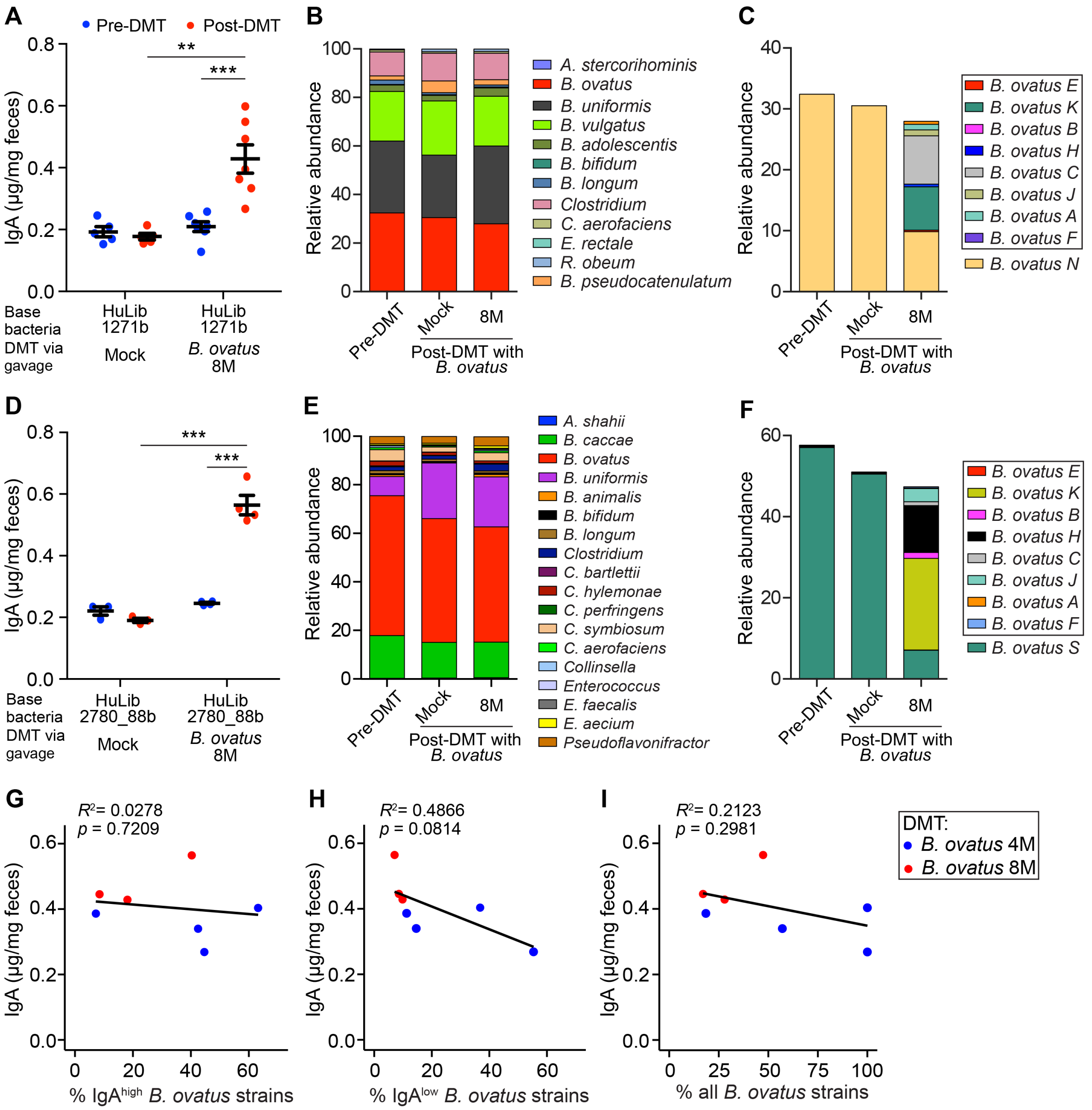


**Figure S10. Robust modification of fecal IgA level with *B. ovatus* cocktails in gnotobiotic mice that were pre-colonized with microbiota arrayed culture collections, related to Figure 4.** (**A-F**) Fecal IgA concentration (**A** and **D**), relative abundance of each bacterial species (**B** and **E**) and relative abundance of different *B. ovatus* strains (**C** and **F**) in gnotobiotic mice pre- and post-DMT. Mice were pre-colonized with microbiota arrayed culture collections (**A**: HuLib1271b; **D**: HuLib2780_88b) for three weeks. Mice were then gavaged with *B. ovatus* 8M. (**G-I**) Correlation between fecal IgA level and relative abundance of IgA$^{high}$ (**G**), IgA$^{low}$ (**H**) and total *B. ovatus* strains (**I**). The averages of fecal IgA level and bacteria relative abundance were used. All mice, being pre-colonized with either single bacterial strain or complex bacterial community for three weeks, were gavaged with either *B. ovatus* 4M or *B. ovatus* 8M. In **A** and **D** plots, data shown are mean ± standard error of the mean and each dot represents a biological replicate. Sequencing plots display the average abundance from three to five mice. Detailed strain information is listed in Tables S4 and S6. *p*-values with statistical significance (assessed by two-tailed Student's *t* test) are indicated: *$p < 0.05$, **$p < 0.01$, ***$p < 0.001$; ns, not significant.

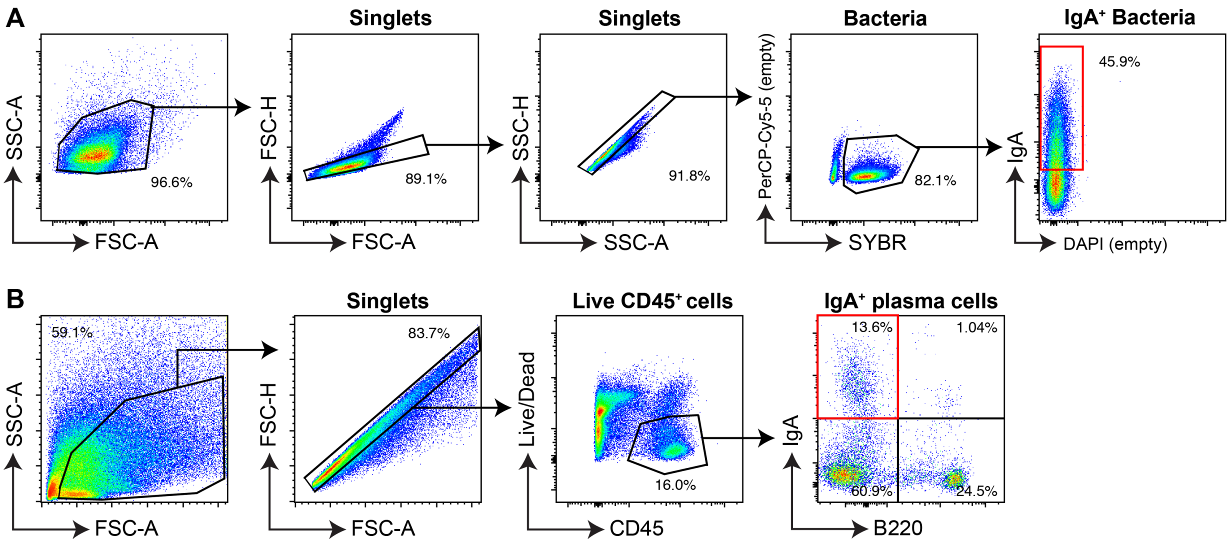


**Figure S11. Representative flow cytometry gating strategies for IgA-coated bacteria and IgA-secreting cells.** (**A**) IgA-coated bacteria in stool were defined as SYBR$^{+}$IgA$^{+}$. (**B**) IgA-secreting B cells in the LP of small intestine and colon were defined as Zombie Aqua$^{-}$ CD45$^{+}$IgA$^{+}$B220$^{-}$.

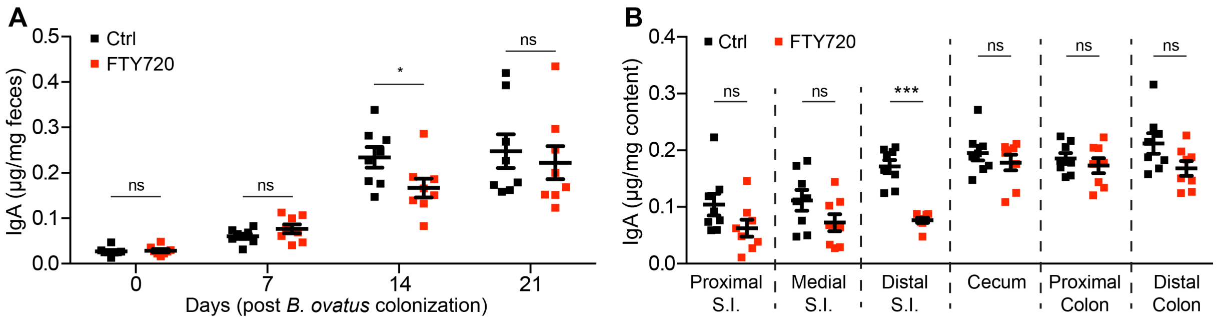


**Figure S12. FTY720 influences luminal IgA production in the distal small intestine but not other regions in mice colonized with *B. ovatus* strain E.** (**A**) Dynamics of fecal IgA concentration in *B. ovatus* strain E colonized gnotobiotic B6 mice treated with or without FTY720. (**B**) Free IgA concentration in different regions along the whole intestinal tract of gnotobiotic mice, which were colonized with *B. ovatus* strains E for three weeks, with or without FTY720 treatment. Data shown are mean ± standard error of the mean. Each dot represents a biological replicate. *p*-values with statistical significance (assessed by unpaired two-tailed Student's *t* test) are indicated: *$p < 0.05$, ***$p < 0.001$; ns, not significant.

### Supplemental tables

**Table S1. Information for each bacterial strain.**

| Phylum | Species | Strain |
|---|---|---|
| Bacteroidetes | *Bacteroides ovatus* | ATCC®8483 |
| Bacteroidetes | *Bacteroides caccae* | ATCC®43185 |
| Bacteroidetes | *Bacteroides thetaiotaomicron* | ATCC®VPI5482 |
| Bacteroidetes | *Bacteroides vulgatus* | ATCC®8482 |
| Firmicutes | *Ruminococcus gnavus* | ATCC®29149 |
| Firmicutes | *Clostridium bolteae* | ATCC®BAA-613 |
| Actinobacteria | *Collinsella aerofaciens* | ATCC® 25986 |
| Proteobacteria | *Escherichia coli* | ATCC®K-12 MG1655 |

**Table S2. Details for different *B. ovatus* strains.**

| Phylum | Species | Strain | Strain Abbreviation |
|---|---|---|---|
| Bacteroidetes | *Bacteroides ovatus* | BSD2780_12_0875_150380_E1 | *B. ovatus A* |
| Bacteroidetes | *Bacteroides ovatus* | 1001095IJ_161003_A6 | *B. ovatus B* |
| Bacteroidetes | *Bacteroides ovatus* | 1001283B150210_160208_F9 | *B. ovatus C* |
| Bacteroidetes | *Bacteroides ovatus* | 1001217B_150727_E1 | *B. ovatus D* |
| Bacteroidetes | *Bacteroides ovatus* | ATCC_8483 | *B. ovatus E* |
| Bacteroidetes | *Bacteroides ovatus* | BSD3178_07_1175_160815_A10 | *B. ovatus F* |
| Bacteroidetes | *Bacteroides ovatus* | BSD3448_08_0949_C3 | *B. ovatus G* |
| Bacteroidetes | *Bacteroides ovatus* | 1001099B_141217_E5 | *B. ovatus H* |
| Bacteroidetes | *Bacteroides ovatus* | 1001275B_160808_G11 | *B. ovatus I* |
| Bacteroidetes | *Bacteroides ovatus* | 1001713B_170207_170306_D4 | *B. ovatus J* |
| Bacteroidetes | *Bacteroides ovatus* | J1101437_171009_F12 | *B. ovatus K* |
| Bacteroidetes | *Bacteroides ovatus* | 1001302B_F3 | *B. ovatus L* |
| Bacteroidetes | *Bacteroides ovatus* | 1001136B_E5 | *B. ovatus M* |
| Bacteroidetes | *Bacteroides ovatus* | 1001271B_150615_H2 | *B. ovatus N* |
| Bacteroidetes | *Bacteroides ovatus* | 1001262B_160229_F6 | *B. ovatus O* |
| Bacteroidetes | *Bacteroides ovatus* | 1001175B_160314_D1 | *B. ovatus P* |
| Bacteroidetes | *Bacteroides ovatus* | BSD2780_06_1687_150420_H2 | *B. ovatus Q* |
| Bacteroidetes | *Bacteroides ovatus* | 1001254J_160919_B1 | *B. ovatus R* |
| Bacteroidetes | *Bacteroides ovatus* | BSD2780_06_1688b_171218_A7 | *B. ovatus S* |

**Table S3. % dissimilarity of genomic DNA sequences amongst various *B. ovatus* strains.**

Different *B. ovatus* strains (columns and rows)

| | D | B | L | K | A | R | O | J | N | P | Q | E | S | F | M | I | C | G | H |
|---|---|---|---|---|---|---|---|---|---|---|---|---|---|---|---|---|---|---|---|
| D | 0.00 | 0.83 | 0.85 | 0.73 | 0.72 | 0.72 | 0.73 | 0.73 | 0.74 | 0.74 | 0.74 | 0.74 | 0.74 | 0.70 | 0.73 | 0.73 | 0.73 | 0.74 | 0.74 |
| B | 0.83 | 0.00 | 0.47 | 0.61 | 0.62 | 0.62 | 0.61 | 0.72 | 0.69 | 0.68 | 0.68 | 0.71 | 0.71 | 0.73 | 0.73 | 0.71 | 0.72 | 0.73 | 0.73 |
| L | 0.85 | 0.47 | 0.00 | 0.67 | 0.67 | 0.63 | 0.64 | 0.68 | 0.69 | 0.70 | 0.70 | 0.71 | 0.70 | 0.73 | 0.71 | 0.72 | 0.73 | 0.73 | 0.72 |
| K | 0.73 | 0.61 | 0.67 | 0.00 | 0.40 | 0.39 | 0.38 | 0.52 | 0.46 | 0.45 | 0.45 | 0.46 | 0.47 | 0.47 | 0.49 | 0.47 | 0.47 | 0.47 | 0.48 |
| A | 0.72 | 0.62 | 0.67 | 0.40 | 0.00 | 0.33 | 0.33 | 0.52 | 0.48 | 0.47 | 0.48 | 0.50 | 0.49 | 0.48 | 0.52 | 0.49 | 0.49 | 0.49 | 0.49 |
| R | 0.72 | 0.62 | 0.63 | 0.39 | 0.33 | 0.00 | 0.03 | 0.50 | 0.46 | 0.47 | 0.47 | 0.50 | 0.48 | 0.48 | 0.52 | 0.50 | 0.51 | 0.51 | 0.52 |
| O | 0.73 | 0.61 | 0.64 | 0.38 | 0.33 | 0.03 | 0.00 | 0.51 | 0.46 | 0.46 | 0.46 | 0.49 | 0.49 | 0.47 | 0.51 | 0.49 | 0.50 | 0.51 | 0.51 |
| J | 0.73 | 0.72 | 0.68 | 0.52 | 0.52 | 0.50 | 0.51 | 0.00 | 0.32 | 0.31 | 0.32 | 0.30 | 0.30 | 0.49 | 0.49 | 0.48 | 0.49 | 0.47 | 0.48 |
| N | 0.74 | 0.69 | 0.69 | 0.46 | 0.48 | 0.46 | 0.46 | 0.32 | 0.00 | 0.09 | 0.09 | 0.27 | 0.29 | 0.46 | 0.42 | 0.40 | 0.43 | 0.43 | 0.44 |
| P | 0.74 | 0.68 | 0.70 | 0.45 | 0.47 | 0.47 | 0.46 | 0.31 | 0.09 | 0.00 | 0.05 | 0.25 | 0.26 | 0.46 | 0.43 | 0.41 | 0.42 | 0.42 | 0.42 |
| Q | 0.74 | 0.68 | 0.70 | 0.45 | 0.48 | 0.47 | 0.46 | 0.32 | 0.09 | 0.05 | 0.00 | 0.25 | 0.27 | 0.46 | 0.43 | 0.41 | 0.42 | 0.42 | 0.43 |
| E | 0.74 | 0.71 | 0.71 | 0.46 | 0.50 | 0.50 | 0.49 | 0.30 | 0.27 | 0.25 | 0.25 | 0.00 | 0.12 | 0.44 | 0.43 | 0.42 | 0.42 | 0.42 | 0.43 |
| S | 0.74 | 0.71 | 0.70 | 0.47 | 0.49 | 0.48 | 0.49 | 0.30 | 0.29 | 0.26 | 0.27 | 0.12 | 0.00 | 0.46 | 0.45 | 0.45 | 0.44 | 0.43 | 0.43 |
| F | 0.70 | 0.73 | 0.73 | 0.47 | 0.48 | 0.48 | 0.47 | 0.49 | 0.46 | 0.46 | 0.46 | 0.44 | 0.46 | 0.00 | 0.40 | 0.41 | 0.41 | 0.40 | 0.41 |
| M | 0.73 | 0.73 | 0.71 | 0.49 | 0.52 | 0.52 | 0.51 | 0.49 | 0.42 | 0.43 | 0.43 | 0.43 | 0.45 | 0.40 | 0.00 | 0.19 | 0.21 | 0.20 | 0.20 |
| I | 0.73 | 0.71 | 0.72 | 0.47 | 0.49 | 0.50 | 0.49 | 0.48 | 0.40 | 0.41 | 0.41 | 0.42 | 0.45 | 0.41 | 0.19 | 0.00 | 0.11 | 0.11 | 0.11 |
| C | 0.73 | 0.72 | 0.73 | 0.47 | 0.49 | 0.51 | 0.50 | 0.49 | 0.43 | 0.42 | 0.42 | 0.42 | 0.44 | 0.41 | 0.21 | 0.11 | 0.00 | 0.11 | 0.10 |
| G | 0.74 | 0.73 | 0.73 | 0.47 | 0.49 | 0.51 | 0.51 | 0.47 | 0.43 | 0.42 | 0.42 | 0.42 | 0.43 | 0.40 | 0.20 | 0.11 | 0.11 | 0.00 | 0.07 |
| H | 0.74 | 0.73 | 0.72 | 0.48 | 0.49 | 0.52 | 0.51 | 0.48 | 0.44 | 0.42 | 0.43 | 0.43 | 0.43 | 0.41 | 0.20 | 0.11 | 0.10 | 0.07 | 0.00 |

**Table S4. Multiplex cocktails of *B. ovatus* strains used in DMT.**

| Cocktail name | Strain |
|---|---|
| *B. ovatus* 4M | *B. ovatus B* |
| | *B. ovatus C* |
| | *B. ovatus E* |
| | *B. ovatus F* |
| *B. ovatus* 8M | *B. ovatus B* |
| | *B. ovatus C* |
| | *B. ovatus E* |
| | *B. ovatus F* |
| | *B. ovatus A* |
| | *B. ovatus H* |
| | *B. ovatus J* |
| | *B. ovatus K* |

**Table S5. Bacterial strains in synthetic cocktail of diverse bacterial species (8member community).**

| Phylum | Species | Strain | Strain Abbreviation |
|---|---|---|---|
| Bacteroidetes | *Bacteroides ovatus* | BSD2780_06_1687_150420_H2 | *B. ovatus Q* |
| Bacteroidetes | *Bacteroides caccae* | ATCC®43185 | *B. caccae* |
| Bacteroidetes | *Bacteroides thetaiotaomicron* | ATCC®VPI5482 | *B. theta.* |
| Bacteroidetes | *Bacteroides vulgatus* | ATCC®8482 | *B. vulgatus* |
| Firmicutes | *Ruminococcus gnavus* | ATCC®29149 | *R. gnavus* |
| Firmicutes | *Clostridium bolteae* | ATCC®BAA-613 | *C. bolteae* |
| Actinobacteria | *Collinsella aerofaciens* | ATCC® 25986 | *C. aero.* |
| Proteobacteria | *Escherichia coli* | ATCC®K-12 MG1655 | *E. coli* |

### Table S6. Bacterial composition in different microbiota arrayed culture collections.

| Library Name | Phylum | Species | Strain | Strain Abbreviation |
|---|---|---|---|---|
| HuLib1175B | Bacteroidetes | *Bacteroides eggerthii* | 1001175st1_B5_1001175B_160314 | *B. eggerthii* |
| | Bacteroidetes | *Bacteroides fragilis* | 1001175st1_C3_1001175B_160314 | *B. fragilis* |
| | Bacteroidetes | *Bacteroides intestinalis* | 1001175st1_A4_1001175B_160314 | *B. intestinalis* |
| | Bacteroidetes | *Bacteroides ovatus* | 1001175st1_E11_1001175B_160314 | *B. ovatus* |
| | Bacteroidetes | *Bacteroides thetaiotaomicron* | 1001175st1_E5_1001175B_160314 | *B. theta.* |
| | Bacteroidetes | *Bacteroides uniformis* | 1001175st1_F6_1001175B_160314 | *B. uniformis* |
| | Bacteroidetes | *Bacteroides vulgatus* | 1001175st1_C6_1001175B_160314 | *B. vulgatus* |
| | Actinobacteria | *Bifidobacterium longum* | 1001175st1_G10_1001175B_160314 | *B. longum* |
| | Firmicutes | *Clostridium 1001175sp1* | 1001175st1_A10_1001175B_160314 | *Clostridium* |
| | Firmicutes | *Clostridium clostridioforme* | 1001175st1_C5_1001175B_160314 | *C. clostridioforme* |
| | Firmicutes | *Clostridium perfringens* | 1001175st1_F9_1001175B_160314 | *C. perfringens* |
| | Firmicutes | *Dorea longicatena* | 1001175st1_H1_1001175B_160314 | *D. longicatena* |
| | Firmicutes | *Enterococcus avium* | 1001175st1_D6_1001175B_160314 | *E. avium* |
| | Proteobacteria | *Escherichia coli* | 1001175st1_F3_1001175B_160314 | *E. coli F3* |
| | Proteobacteria | *Escherichia coli* | 1001175st2_F4_1001175B_160314 | *E. coli F4* |
| | Proteobacteria | *Escherichia coli* | 1001175st3_A2_1001175B_160314 | *E. coli A2* |
| | Firmicutes | *Lachnospiraceae 1001136sp1* | 1001175st1_C9_1001175B_160314 | *Lachnospiraceae* |
| | Firmicutes | *Lactobacillus 1001175sp1* | 1001175st1_D8_1001175B_160314 | *Lactobacillus* |
| | Bacteroidetes | *Parabacteroides merdae* | 1001175st1_A1_1001175B_160314 | *P. merdae* |
| | Firmicutes | *Roseburia 1001271sp1* | 1001175st1_E3_1001175B_160314 | *Roseburia* |
| | Firmicutes | *Ruminococcus 1001175sp1* | 1001175st1_E1_1001175B_160314 | *Ruminococcus* |
| | Firmicutes | *Streptococcus 1001175sp1* | 1001175st1_H6_1001175B_160314 | *Streptococcus H6* |
| | Firmicutes | *Streptococcus 1001283sp2* | 1001175st1_H3_1001175B_160314 | *Streptococcus H3* |
| | Firmicutes | *Streptococcus anginosus* | 1001175st1_H11_1001175B_160314 | *S. anginosus* |
| HuLib1271b | Firmicutes | *Anaerofustis stercorihominis* | 1001271st1_D3_1001271B_150615 | *A. stercorihominis* |
| | Bacteroidetes | *Bacteroides ovatus* | 1001271st1_H2_1001271B_150615 | *B. ovatus* |
| | Bacteroidetes | *Bacteroides uniformis* | 1001271st1_A10_1001271B_150615 | *B. uniformis* |
| | Bacteroidetes | *Bacteroides vulgatus* | 1001271st1_G7_1001271B_150615 | *B. vulgatus* |
| | Actinobacteria | *Bifidobacterium adolescentis* | 1001271st1_A4_1001271B_150615 | *B. adolescentis* |
| | Actinobacteria | *Bifidobacterium bifidum* | 1001271st1_H11_1001271B_150615 | *B. bifidum* |
| | Actinobacteria | *Bifidobacterium longum* | 1001271st1_B4_1001271B_150615 | *B. longum* |
| | Actinobacteria | *Bifidobacterium pseudocatenulatum* | 1001271st1_F3_1001271B_150615 | *B. pseudocatenulatum* |
| | Firmicutes | *Clostridium 1001271sp1* | 1001271st1_H5_1001271B_150615 | *Clostridium* |
| | Actinobacteria | *Collinsella aerofaciens* | 1001271st1_C3_1001271B_150615 | *C. aerofaciens* |
| | Firmicutes | *Eubacterium rectale* | 1001271st1_F12_1001271B_150615 | *E. rectale* |
| | Firmicutes | *Ruminococcus obeum* | 1001271st1_E5_1001271B_150615 | *R. obeum* |
| HuLib2780_88b | Bacteroidetes | *Alistipes shahii* | BSD2780061688st1_A10_BSD2780061688b_171218 | *A. shahii* |
| | Bacteroidetes | *Bacteroides caccae* | BSD2780061689st1_A4_BSD2780061688b_171218 | *B. caccae* |
| | Bacteroidetes | *Bacteroides ovatus* | BSD2780061688st1_C6_BSD2780061688b_171218 | *B. ovatus* |
| | Bacteroidetes | *Bacteroides uniformis* | BSD2780061689st1_G7_BSD2780061688b_171218 | *B. uniformis* |
| | Actinobacteria | *Bifidobacterium animalis* | BSD2780061688st1_E5_BSD2780061688b_171218 | *B. animalis* |
| | Actinobacteria | *Bifidobacterium bifidum* | BSD2780061688st1_G1_BSD2780061688b_171218 | *B. bifidum* |
| | Actinobacteria | *Bifidobacterium longum* | BSD2780061688st2_H1_BSD2780061688b_171218 | *B. longum* |
| | Firmicutes | *Clostridium BSD2780061688sp2* | BSD2780061688st1_H5_BSD2780061688b_171218 | *Clostridium E5* |
| | Firmicutes | *Clostridium BSD2780061688sp3* | BSD2780061688st1_E8_BSD2780061688b_171218 | *Clostridium E8* |
| | Firmicutes | *Clostridium bartlettii* | BSD2780061688st1_A9_BSD2780061688b_171218 | *C. bartlettii* |
| | Firmicutes | *Clostridium hylemonae* | BSD2780061688st1_A6_BSD2780061688b_171218 | *C. hylemonae* |
| | Firmicutes | *Clostridium perfringens* | BSD2780061688st3_G3_BSD2780061688b_171218 | *C. perfringens* |
| | Firmicutes | *Clostridium symbiosum* | BSD2780061688st1_G6_BSD2780061688b_171218 | *C. symbiosum* |
| | Actinobacteria | *Collinsella aerofaciens* | BSD2780061688st1_F5_BSD2780061688b_171218 | *C. aerofaciens* |
| | Actinobacteria | *Collinsella species* | BSD2780061688st1_H8_BSD2780061688b_171218 | *C. species* |
| | Firmicutes | *Enterococcus 1001136sp1* | BSD2780061688st2_D3_BSD2780061688b_171218 | *Enterococcus* |
| | Firmicutes | *Enterococcus faecalis* | BSD2780061688st3_G10_BSD2780061688b_171218 | *E. faecalis* |
| | Firmicutes | *Enterococcus faecium* | BSD2780061688st2_C8_BSD2780061688b_171218 | *E. faecium* |
| | Firmicutes | *Pseudoflavonifractor BSD2780061688sp1* | BSD2780061688st1_E11_BSD2780061688b_171218 | *Pseudoflavonifractor* |

**Table S7. Detailed information about various bacterial strains.**

| Phylum | Species | Strain | Strain Abbreviation |
|---|---|---|---|
| Bacteroidetes | *Bacteroides ovatus* | ATCC®8483 | *B. ovatus E* |
| Bacteroidetes | *Bacteroides caccae* | ATCC®43185 | *B. caccae A* |
| Bacteroidetes | *Bacteroides caccae* | 1001285I_161205_F12 | *B. caccae B* |
| Bacteroidetes | *Bacteroides caccae* | BSD3178_07_1176_160815_A7 | *B. caccae C* |
| Bacteroidetes | *Bacteroides thetaiotaomicron* | ATCC®VPI5482 | *B. theta. A* |
| Bacteroidetes | *Bacteroides thetaiotaomicron* | BSD2780_12_0875b_A6 | *B. theta. B* |
| Bacteroidetes | *Bacteroides thetaiotaomicron* | BSD2780_06_1689_150309_F9 | *B. theta. C* |
| Bacteroidetes | *Bacteroides vulgatus* | ATCC®8482 | *B. vulgatus A* |
| Bacteroidetes | *Bacteroides vulgatus* | BSD2780_12_0874b_170522_A7 | *B. vulgatus B* |
| Bacteroidetes | *Bacteroides vulgatus* | 1001271B_150615_G7 | *B. vulgatus* C |
| Bacteroidetes | *Parabacteroides johnsonii* | DSMZ_18315 | *P. johnsonii* |
| Bacteroidetes | *Bacteroides intestinalis* | DSMZ_17393 | *B. intestinalis* |
| Bacteroidetes | *Bacteroides fragilis* | J1001437_171009_C3 | *B. fragilis* |
